# A covarion model for phylogenetic estimation using discrete morphological datasets

**DOI:** 10.1101/2025.06.20.660793

**Authors:** Basanta Khakurel, Sebastian Höhna

## Abstract

The rate of evolution of a single morphological character is not homogeneous across the phylogeny and this rate heterogeneity varies between morphological characters. However, traditional models of morphological character evolution often assume that all characters evolve according to a time-homogeneous Markov process, which applies uniformly across the entire phylogeny. While models incorporating among-character rate variation alleviate the assumption of the same rate for all characters, they still fail to address lineage-specific rate variation for individual characters. The covarion model, originally developed for molecular data to model the invariability of some sites for parts of the phylogeny, provides a promising framework for addressing this issue in morphological phylogenetics. In this study, we extend the covarion model in `RevBayes` to morphological character evolution, which we call the *covariomorph* model, and apply it to a diverse range of morphological datasets. Our covariomorph model utilizes multiple rate categories derived from a discretized probability distribution, which scales rate matrices accordingly. Characters are allowed to evolve within any of these rate categories, with the possibility of switching between rate categories during the evolutionary process. We verified our implementation of the covariomorph model with the help of simulations. Additionally, we examined 164 empirical datasets, finding patterns of rate heterogeneity compatible with covarion-like dynamics in approximately half of them. Upon further examination of two focal datasets that exhibited covarion-like rate variation, we found that the covariomorph model provides a more nuanced approach to incorporate rate variation across lineages, significantly affecting the resulting tree topology and branch lengths compared to traditional models. The observed sensitivity of branch lengths to model choice underscores potential implications of this approach for divergence time estimation and evolutionary rate calculations. By accounting for lineage- and character-specific rate shifts, the covariomorph model offers a robust framework to improve the accuracy of morphological phylogenetic inference.

## 1 Introduction

Phylogenetic inferences from morphological character data using likelihood-based methods commonly employ the Markov *k*-state (Mk) model (Lewis, 2001). The Mk model represents a generalization of the Jukes-Cantor (Jukes and Cantor, 1969) model of molecular evolution, extending its principles to morphological data. Under the Mk model, characters in a morphological data matrix are assumed to have equal transition rates among character states. Furthermore, the Mk model assumes equal rates across all branches of the phylogeny (when the branch lengths are represented in expected number of transitions between character states). However, the Mk model may not fit the biological realism of morphological evolution due to its simplistic assumptions (Wright et al., 2016; Lee and Palci, 2015; O’Reilly et al., 2018). In particular, the assumption of homogeneous rates of evolution for all characters across the entire phylogeny may be violated, as selective constraints on a set of characters due to ecological shifts can vary between lineages over time. (Skinner, 2010; Wang and Lloyd, 2016; Lloyd et al., 2012).

A compelling example of rate heterogeneity in morphological evolution across lineages can be found in early avian evolution. Wang and Lloyd (2016) demonstrated that, during early avian evolution, different anatomical regions evolved at differential rates as adaptations for flight emerged. For instance, the rapid modification of forelimbs occurred concurrently with skeletal structures supporting wings, while other anatomical features evolved more gradually. Furthermore, subsequent ecological specialization across avian lineages has led to variable rates of character evolution across different anatomical regions, reflecting diverse selective pressures in different niches. These types of pattern where different character complexes evolve at dissimilar rates within a group of organism has been documented across diverse taxonomic groups, from arthropods to vertebrates, suggesting that models assuming uniform rates for characters across branches may be fundamentally misspecified for many empirical datasets (Clarke and Middleton, 2008; Skinner, 2010; Brocklehurst, 2017).

To relax the assumptions of the Mk model and account for rate heterogeneity, various approaches have been proposed. A schematic showing different approaches for modeling rate variation can be seen in Fig. 1. One widely used method to incorporate among character rate variation (ACRV) is to assume that each character evolves under one of the *r* discrete rate categories obtained from the quantiles of a gamma distribution (+Γ; Fig. 1 middle row; Yang, 1994; Capobianco and Hö hna, 2025). Additionally, some studies have suggested the use of a lognormal distribution as an alternative and concluded that incorporating among character rate variation is vital for robust phylogenetic inferences (Wagner, 2012; Harrison and Larsson, 2015). However, a limitation of such among character rate variation model is that the fast-evolving characters are always fast and the slow-evolving characters are always slow, which might not be biologically realistic.

**Figure 1.**
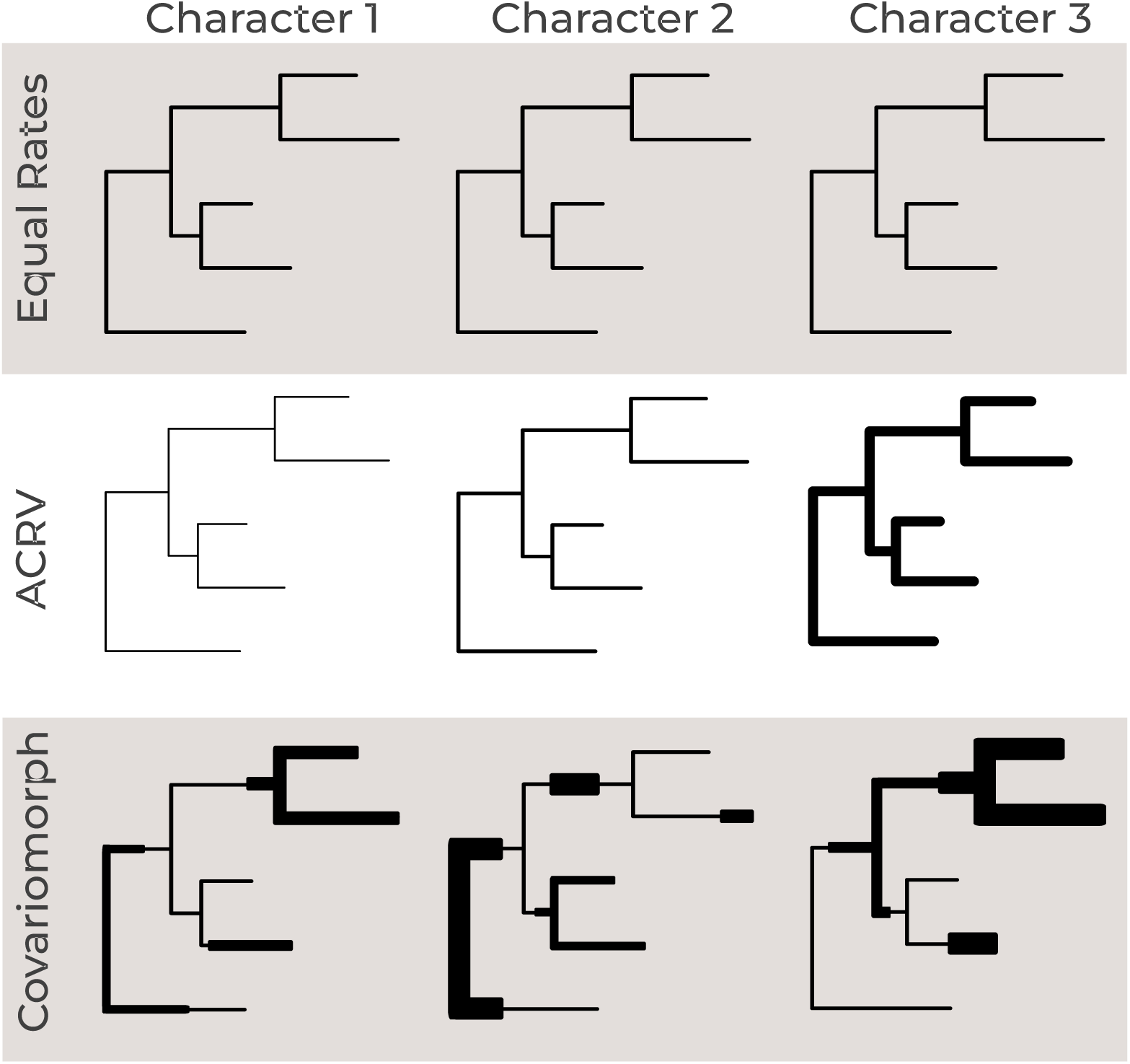
A schematic illustrating different approaches for modeling evolutionary rates in morphological phylogenetics. We demonstrate these models for three characters across the phylogeny to highlight the heterogeneity across characters and across lineages. The top row depicts the equal rates model (i.e., standard Mk model; Lewis, 2001), where all branches maintain uniform evolutionary rates across characters and lineages. The middle row represents the among-character rate variation (ACRV) model, highlighting rate variation across characters but homogeneous across lineages. For example, character 3 shows accelerated rates of evolution (indicated by thicker branches) while character 1 shows decelerated rates of evolution. The bottom row demonstrates the covariomorph model, where evolutionary rates vary along lineage, as shown by the varying branch thicknesses within each character tree. The covariomorph model implicitly includes rate variation among character because each character has its own lineage-specific rate variation. This visualization effectively contrasts how these three models conceptualize different patterns of evolutionary rate heterogeneity. Figure inspired from Galtier (2001).

Importantly, ACRV addresses heterogeneity among characters, but not heterogeneity among branches. Relaxed morphological clocks provide a complementary approach by allowing branch-specific rate multipliers but those rate multipliers apply uniformly to all characters (Beck and Lee, 2014; Lee et al., 2014). Effectively, a morphological relaxed clock model functions similarly to an unrooted phylogeny with branch lengths measured in expected number of transitions where long branches represent fast rates with many expected changes. This approach, while useful, imposes the significant constraint that a branch exhibiting a faster rate of evolution is necessarily fast for all characters; an assumption that may be violated in morphological datasets.

A more nuanced approach to modeling morphological evolution involves incorporating branch length variation among sites, a phenomenon known as heterotachy (Lopez et al., 2002; Zhou et al., 2007). Heterotachy can be addressed through several approaches. First, a commonly used approach is data partitioning, where datasets are divided into subsets based on *a priori* assumptions about character evolution, allowing each subset to evolve under different model parameters. For example, a dataset can be partitioned by anatomical region (Casali et al., 2023) or by homoplasy (Rosa et al., 2019). Furthermore, datasets can be partitioned by size of the state space per character, e.g., creating a data subset for all binary characters, another data subset for all 3-state characters, and so forth (Khakurel et al., 2024). Approaches to partitioning a morphological matrix might relax the assumptions that all characters have the same evolutionary rate throughout the phylogeny but do not necessarily accommodate lineage-specific rate variation within each character (except if each character would be placed into its own data subset). Additionally, data partitioning is not straightforward and the optimal partitioning strategy is not always obvious with different schemes producing conflicting results (Casali et al., 2022).

An alternative to dataset partitioning is explicitly modeling heterotachy (Fig. 1, bottom row). Modeling heterotachy has been attempted using sets of branch lengths to set up a mixture model (Meade and Pagel, 2008; Crotty et al., 2020). Alternatively, the number of branch length sets can be a part of the analyses, e.g., by employing reversible-jump Markov chain Monte Carlo algorithm (rjMCMC) across the sets of branches (Pagel and Meade, 2008). These mixture model approaches are effectively data partitioning approaches where the assignment to data subsets and possibly the number of data subsets is not specified *a priori*. Besides these phenomenological models, heterotachy has also been modeled explicitly by utilizing a more mechanistic model: the covarion model (Tuffley and Steel, 1998; Galtier, 2001; Huelsenbeck, 2002; Toups et al., 2024). The covarion model, originally described by Fitch and Markowitz (1970) and implemented by Tuffley and Steel (1998) for molecular data, accounts for sites that at some lineages have very low (or zero) mutation rates and at other lineages have the ability to mutate. This covarion model allows for sites to exist in either “on” or “off” state along a phylogeny which thus results in site-specific rate variation along lineages (Tuffley and Steel, 1998; Penny et al., 2001; Shavit Grievink et al., 2008). In addition to the “off” and “on” covarion model, other studies have explored extensions of the original covarion model (Galtier, 2001; Penny et al., 2001; Pupko and Galtier, 2002). These studies refer to the covarion model as a more general model in which sites may undergo some switching between different rate regimes as characters evolve along the tree.

In morphological phylogenetics, however, the idea of using covarion-like transition models has not been tested previously. The prevailing impact of differential selection pressures on differently evolving characters lead us to speculate that the evolution of characters along a phylogenetic tree might not be as uniform as previously assumed in the simpler models. Furthermore, Goloboff et al. (2019) have shown, with both empirical and simulated data, that morphological datasets refute the idea of common branch lengths for all characters. Thus, a ‘common mechanism’ model such as the Mk and its additional extensions (e.g., Mkv or mMkv+Γ; Lewis, 2001; Capobianco and Hö hna, 2025) do not provide a sufficient fit for morphological evolution and a model incorporating heterotachy could more accurately represent the nature of morphological datasets.

In this study, we develop and apply a modified covarion model to allow for rates of morphological evolution for different characters to vary over lineages in morphological phylogenetic datasets. We call our new model the *“covariomorph”* model, following the spirit of the covariotide model for nucleotides (Lockhart et al., 1998; Huelsenbeck, 2002). Our covariomorph model extends the approach of the original covarion model to accommodate multiple rate categories, thus allowing for a much wider range of rate variation (Fig. 1). This newly proposed covariomorph model relaxes the restrictive assumption of homogeneous character evolution rate across the phylogeny. We explore our covariomorph approach by means of simulations specifically focusing at the switching between rate regimes. We finally apply the covariomorph model to a range of published empirical datasets and explore its utility in phylogenetic tree estimation using morphological datasets. As we show, a notable amount of empirical datasets exhibit switching between different rate regimes, resulting in significant differences in branch lengths of the resulting phylogeny, thus highlighting the utility of our covariomorph model.

## 2 Methods

In general, our *covariomorph* approach is similar to approaches proposed by Galtier (2001) and Penny et al. (2001) where they assume that for each character there are two stochastic processes acting upon it; one making changes among a number of rates of evolution, and the other making changes according to the current rate of evolution among a number of character states.

### 2.1 The Covariomorph Process

Let us begin by defining the general *covariomorph* process. As an illustration, we consider a single character, and characters in a data matrix are assumed to be independent and identically distributed. The covariomorph process is a continuous time Markov process along a phylogeny. The process initiates by drawing a rate category *r* for the character from a specified prior probability distribution, which may be either discrete or continuous. Concurrently, the character’s initial state is drawn from a separate prior distribution, for example, a uniform distribution where each state is equally probable *a priori*.

As the character evolves, its state transitions are governed by a base substitution rate matrix *Q* (in our case, the Mk model), scaled by the current rate *r*. The total rate of leaving any given state *i* is −*rQ_ii_* where *Q_ii_* is the diagonal element of the substitution rate matrix corresponding to state *i*. In parallel, the rate category *r* itself can change in a rate switching event, which occurs at a constant rate *δ*.

Therefore, as the character evolves along a branch of the phylogeny, two types of stochastic events can occur. The waiting time for any event is governed by an exponential distribution with a total rate equal to the sum of the rates of both event types, *δ* − *rQ_ii_*:

- With probability 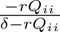, a state transition event occurs changing the observable state of the character (e.g., from state 0 to state 1). The specific rate of transitions between observable states is determined by a selected substitution model *rQ*.
- With probability 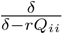, a rate shift event occurs changing the underlying rate itself. A new rate category is drawn from the prior distribution. This new rate subsequently modulates the probability of future state transitions, i.e., rescales the transition matrix of the Mk model. Consequently, a character evolving at a certain pace can instantaneously switch to a faster or slower rate at any point along a branch.

At each internal node (speciation event), the current character state and rate category are inherited by both descendant lineages (symmetric inheritance, no cladogenetic events). The evolutionary process then continues independently along each daughter branch. This entire simulation of drawing an initial rate, simulating transitions and rate shifts along branches, and propagating states and rates at nodes is repeated until the terminal nodes (tips) of the phylogeny are reached. This yields the final character states, which represents the observed data, while rate categories remain latent. The complete process is then performed independently for each morphological character in the dataset.

### 2.2 The covariomorph model

Our *covariomorph* model is an explicit case of the general covariomorph process described above, with a fixed number of possible rate categories (i.e., a discrete prior distribution on the number of rates). We construct our covariomorph model using several simple extensions to the Mk model rate matrix (Q-matrix). Our covariomorph model consists of three components: (i) an underlying model of discrete character evolution with *k*-states, which we assume here as the Mk model (Lewis, 2001), (ii) a set **r** of *m* number of rates to rescale the *k*-state rate matrix *Q*, with condition 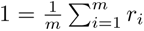 so that the rates are normalized, and (iii) a switching rate *δ* between the rate categories.

The covariomorph rate matrix Q̃ is constructed as

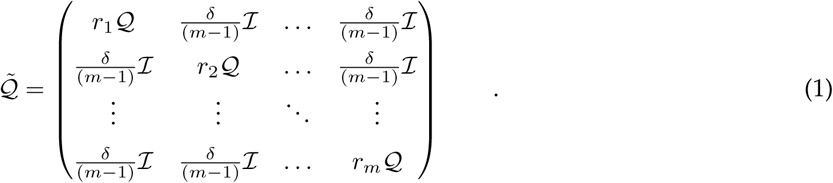

where I is an identity matrix of size *k*. Note that the original character matrix Q has size *k* while our covariomorph rate matrix Q̃ has size *k* × *m*. Thus, the resulting covariomorph rate matrix Q̃ effectively expands the state space to account for both the character states and their associated rate categories. The diagonal blocks (*r_i_*Q) represent the evolutionary processes occurring within each rate category, while the off-diagonal blocks 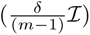 capture the rate shift events from one rate category to another.

#### 2.2.1 Per-category rate scalars

In our covariomorph model, the rate scalars (**r**) are obtained from a discretized probability distribution. For example, analogous to the modeling of among-character rate variation in phylogenetic analyses, we can use a gamma (Yang, 1994) or a lognormal (Harrison and Larsson, 2015) distribution to obtain the rate scalars. Each rate scalar is then used to scale the Mk model’s Q-matrix, effectively generating a set of scaled rate matrices. This scaling permits different rate categories to exhibit different transition dynamics, allowing characters to evolve at variable rates depending on the assigned rate category. In our examples, we will use a lognormal distribution with median parameter *η* = 1 in real-space and standard deviation *σ* determining the scale of rate variation. The rates are then computed as the medians of the *m* quantiles

and normalized afterwards to sum to one, 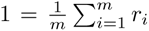. The normalization of rate scalars allows us to interpret branch lengths in our covariomorph model as the expected number of character state transitions per character. This is because switching between rate categories changes only the evolutionary rate regime, not the character state itself.

#### 2.2.2 Switching between rate categories

We define a switching rate parameter *δ* that governs the process to switch between rate categories, and is assumed to be homogeneous among all rate categories. Thus, for simplicity of the model, we assume a symmetric switching mechanism where the rate of transitioning between rate categories is uniform, meaning the probability of moving from one rate category to another is the same regardless of the current category. The switching process only permits changes in the rate regime, ensuring that the substitution process within the character states is governed solely by the scaled Mk matrices. Furthermore, we re-parameterize the switching rate by dividing it with the total number of rate categories minus one 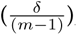, which normalizes the switching rate across all possible rate category transitions. This reparameterization allows the switching rate to be comparable across different number of rate categories *m*.

#### 2.2.3 Example 3-rate binary covariomorph model

As an example, let us consider a covariomorph rate matrix with three rate categories for a binary morphological character (Fig. 2). The Q-matrix of a binary character under the Mk model can be represented by a 2 × 2 matrix:

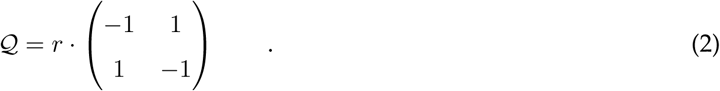

**Figure 2.**
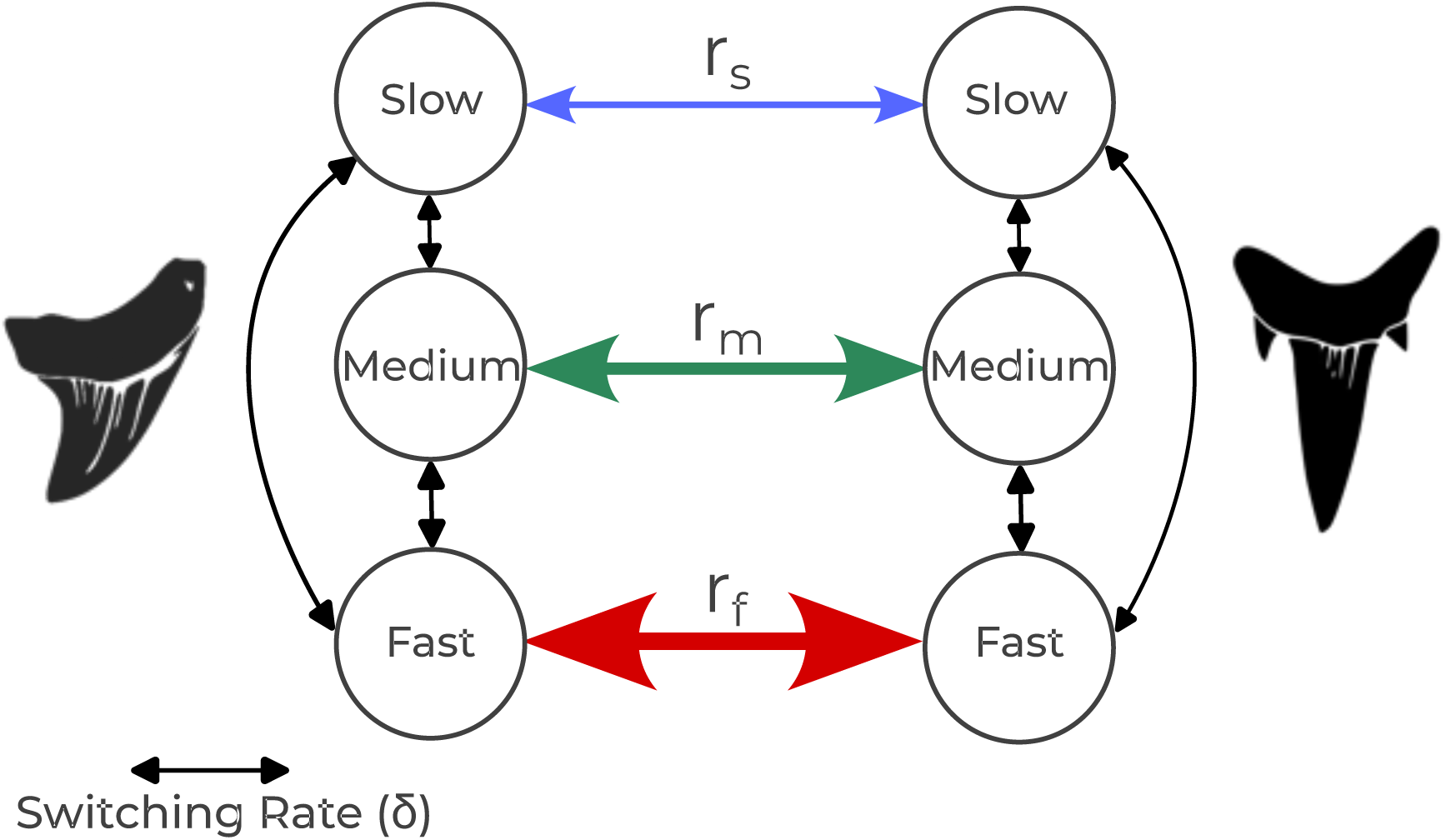
A schematic showing of the 3-rate binary covariomorph model. A character (represented here as different tooth shapes) can be in one of two states and evolve under one of the three rates of evolution: slow(r_s_), medium (r_m_), or fast (r_f_). State transitions occur at one of these rates (colored arrows). The characters can also switch between rate categories (e.g., from medium to fast) at a rate governed by δ (black arrows). This process allows evolutionary rates to vary among characters and across lineages.

where *r* is a scaling factor introduced to scale the matrix such that the branch lengths can be interpreted in units of expected number of character transitions per unit of time.

##### Switching Rate (δ)

To allow for rate heterogeneity, the covariomorph model expands on this by introducing different rate categories. In this example, we use three categories: slow, medium, and fast, defined by rate scalars *r_s_, r_m_* and *r_f_* (see Fig. 2). The rate scalars are derived from a normalized discrete probability distribution such that their mean is 1 (i.e., (*r_s_* + *r_m_* + *r_f_*)*/*3 = 1). Each rate category thus represents a different evolutionary tempo, allowing some characters to evolve faster or slower than others.

A switching rate parameter, *δ*, controls the rate of transition between these rate categories. Combining the character state transitions and the rate switching process, we can construct a 6 × 6 covariomorph Q-matrix (Q̃):

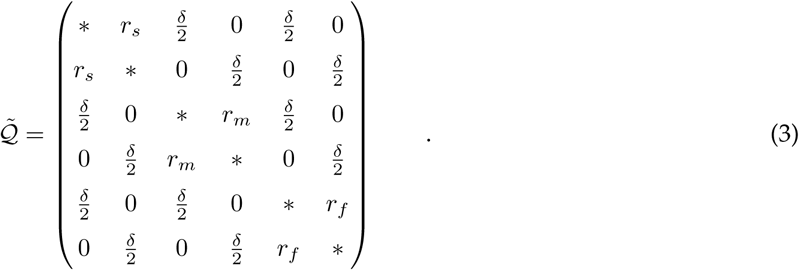

In this rate matrix:

- the rates *r_s_, r_m_* and *r_f_* govern transitions between character state 0 and 1 within a rate category.
- the rate *δ/*2 represents the transition between rate categories (e.g., from slow to medium), assuming an equal probability of switching to either of the other two categories.
- the diagonal elements (∗) are negative values calculated so that each row sums to zero (i.e., Q̃_*ii*_ = — Σ_j≠i_ Q̃_ij_).

#### 2.2.4 State space and likelihood calculation

The covariomorph rate matrix Q̃ introduces *virtual* states. Each virtual state corresponds to a specific character state observed in the data matrix (0 or 1 in case of a binary character) combined with a particular rate category. This means that instead of just *k* states, the model now considers multiple versions of each state each associated with a different evolutionary rate categories. For example, in a 3-rate binary model, the data matrix is expanded into its virtual states and the original state 0 is represented in the expanded matrix by [0, 2, 4] whereas the original state 1 is represented by [1, 3, 5] (Supplementary Fig. S3). This expansion systematically transforms each character of the original morphological matrix into ambiguous virtual characters. Thus, the partial likelihood vectors at the tip of a phylogeny are initialized with 1 for states *i*(2× 1)*, i*(2× 2)*, . . ., i*(2× *m*) when the actually observed state is *i* and initialized as 0 otherwise, while an unobserved state (missing data) is initialized with 1s for all virtual states. This initialization at the tips provides the starting conditions for likelihood calculations, which then proceed recursively through the tree using Felsenstein’s pruning algorithm (Felsenstein, 1981). The pruning algorithm then marginalizes over all possible virtual states of the characters in the internal nodes. Thus, any standard phylogenetic likelihood machinery can be used if implemented for appropriate state space (Smith et al., 2024).

#### 2.2.5 Root frequencies of the covariomorph model

For our covariomorph model, the prior distribution at the root is structured with equal probabilities across both dimensions. We treat all rate categories and character states as equally likely at the root of the phylogeny. First, for the rate categories, each character has an equal probability (1*/m*) of belonging to any of the *m* rate categories. Second, within each rate category, the character state follows the standard Mk model assumption, where each of the *k* possible character state has an equal probability (1*/k*) of occurring. This creates a uniform joint prior distribution across the entire expanded state space (*m* × *k* dimensions), where the probability of any specific combination of rate category and character state is 1*/*(*m* × *k*). For example, in a model with 3 rate categories and binary character states, each of the 6 possible combinations would have a prior probability of 1/6 at the root.

#### 2.2.6 Theoretical properties and relationship to the Mk and Mk+ACRV models

To clarify the relationship of the covariomorph model with existing frameworks, it is instructive to consider the theoretical behavior of the covariomorph model under several limiting conditions. As a generalization of the Mk and Mk+ACRV models, its behavior is expected to converge to these simpler models when certain parameters are pushed to their extremes.

First, consider a scenario where the rate-switching parameter approaches zero (*δ* → 0). When the rate categories differ significantly (*σ >* 0) but switching between them is absent (*δ* ≈ 0), then the model collapses to an Mk model with among character rate variation. The connection is that our covariomorph with distinctive rate categories *m* is equivalent to the ACRV model with rate categories *m* if no switching between the rate categories along the phylogeny is allowed.

Conversely, the model’s behavior is also expected to collapse to that of the Mk model under two other conditions.

- When all rate categories are equal (*σ* ≈ 0), then switching becomes meaningless, as moving between identical rates has no meaningful effect. In this case, the model collapses into the Mk model, with the switching rate effectively reflecting the prior mean rather than any real biological process.
- A less intuitive prediction is that the covariomorph model should also converge to an Mk model if the switching rate is extremely high (e.g., *δ >* 5, which means that 5-times as many rate switches occur compared with actual character state transitions). The expectation is that with very frequent switching between rate categories minimizes the variability in rates across lineages, effectively blending them into a uniform pattern. Under these conditions, the high rate of switching overrides any distinctive rate differences, causing the model to converge into the Mk model. The inference of an elevated switching rate suggests that any detectable rate variation is overshadowed by rapid rate transitions. These frequent transitions could be interpreted as an artifact of the model or a reflection of model constrains rather than representing a true biological process.

Understanding these theoretical boundaries is crucial for interpreting the model’s inferences. We test these behaviors of the covariomorph model via simulations (see sections Methods: Simulation Study and Results: Simulation Results). When the model infers parameters approaching these limits, our findings suggest that the data lack a strong signal for the more complex dynamics of the full covariomorph process.

#### 2.2.7 Implementation

We implemented and validated our covariomorph model within the Bayesian phylogenetic software `RevBayes` (Hö hna et al., 2016), which provides a flexible and extensible framework for probabilistic modeling (Hö hna et al., 2014, Supplementary Figs. S1, S2). In `RevBayes`, the *expandCharacters()* function allows users to expand the original data matrix according to a user-specified number of rate categories. Additionally, to construct the covariomorph rate matrix, the function *fnCovarion()* integrates discrete rate categories with a switching process between them. This function takes in three key arguments: (i) `RateScalars`, defining the relative rate multipliers for each rate category; (ii) `SwitchRates`, specifying the transition rates between the *m* rate categories; and (iii) `RateMatrices`, specifying the rate matrices of the base substitution model that is rescaled. An example `RevBayes` script showing the use of this model can be found in the online supplementary material (available in Dryad).

The covariomorph model’s expanded state space (*k* × *m*) and additional parameters (a switching rate *δ*, and a rate variation parameter *σ*) necessarily increase computational cost compared to the standard Mk or Mk+ACRV models. The computational runtime increases quadratically with the number of rate categories (*m*), and memory requirements scale linearly with the number of taxa, patterns, states, and rate categories. However, our benchmarks show that while the covariomorph model is more computationally intensive, it remains practical for phylogenetic analyses (Supplementary Fig. S4).

### 2.3 Simulation Study

Our simulation study focused on evaluating the model’s ability to recover the true switching rates and detect phylogenetic signals indicative of covarion-like rate variation. To comprehensively evaluate the model’s behavior under diverse evolutionary scenarios, we performed three sets of simulation using varying tree sizes and tree lengths. We simulated datasets on three empirical phylogenies with varying total tree lengths (TL), measured in expected number of transitions per character. The phylogenies selected were:

- A short tree with a tree length of 3.18 and 21 taxa derived from a study of extinct and extant tortoises (Valenti et al., 2022).
- A intermediate length empirical tree of vascular plants (Euphyllophytes) consisting of 56 taxa and a tree length of 10.54 (Rothwell and Nixon, 2006).
- A long tree with 121 taxa and a tree length of 16.71 obtained from a study of cretaceous mammals (Mao et al., 2021).

The total tree length is a crucial factor, as it provides an idea of the expected number of switches between rate categories. For example, for a tree length of 10.54, a switching rate of *δ* = 0.5, corresponds to an average of 5.3 expected switches between rate categories per character across the tree. For each tree length, we generated 100 simulated datasets each for eight different switching rates *δ* ∈ {0.005, 0.05, 0.5, 1, 2, 3, 4, 5}

using our four-rate covariomorph model. For the base rate matrix, we used an observed state space of four to simulate characters. In this case, it is still possible to simulate binary or 3-state characters in addition to 4-state characters. Apart from the scenarios involving switching, we investigated two scenarios representing conditions with no switching between the rate categories (i.e., *δ* = 0). The first scenario assumed characters evolving under a standard Mk model (all characters share a uniform evolutionary rate), and the second followed an Mk + ACRV model (characters have varying rates but constant across the phylogeny). The framework for among-character rate variation model was set similar to the covarion-like rate variation (see below).

To introduce covarion-like rate variation in the simulated datasets, we used a discretized lognormal distribution for rate scalars (Supplementary Fig. S5). We specified the distribution’s mean parameter to 0 (i.e., arithmetic median *η* = 1), which allows us to use the standard deviation to control the shape and range of rate variation. A standard deviation of *σ* = 1.174 was chosen for the simulation of datasets. This value was obtained by first calculating the standard deviation required to span one order of magnitude across a 95% probability interval and then doubling that value, producing a 95% probability interval spanning two orders of magnitude (Hö hna et al., 2017).

We utilized the simulation functionalities within `RevBayes` (e.g., the phylogenetic continuous-time Markov chain (CTMC) process implementation), in order to simulate datasets under our covariomorph model. This simulation process operates directly on the expanded virtual states, meaning that the raw simulated datasets include labels for these virtual states. However, because empirical data matrices typically do not retain virtual state information, we applied a custom R script (available in Dryad) to transform the simulated data matrices, making them resemble the empirical data matrices.

For the inference of simulated datasets, we used a 4-rate covariomorph model with selected prior distributions to prevent excessive constraints on rate variation and switching, ensuring that the posterior estimates are primarily informed by the data rather than strong prior assumptions. We did not partition the dataset by maximum number of states as the datasets were generated using an unpartitioned rate matrix (using an observed state space of 4). The analyses required specifications for prior distributions for the two newly introduced parameters: the standard deviation *σ* of the lognormal distribution to obtain a set of rate scalars and the switching rate *δ*. The rate scalars were derived from a discretized lognormal distribution with arithmetic median of 1 and a prior on the inverse of the standard deviation set as a uniform distribution from 0 to 10^3^

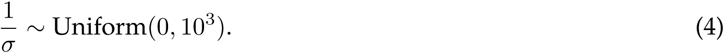

We specified this prior distribution to induce less *a priori* expected rate variation between the rate categories. Similarly, the prior on switching rate (*δ*) was specified as a uniform distribution from 0 to 10^4^

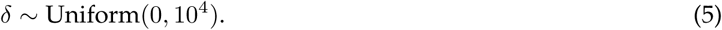

The model assumed a constant switching rate between the rate categories to reduce model complexity.

We assumed a uniform prior on the unrooted tree topologies with branch lengths obtained from independent exponential distributions each with a mean of 0.1.

We performed Markov chain Monte Carlo (MCMC) analyses with four independent replicates for 100,000 generations each, with convergence assessed using `convenience` (Fabreti and Hö hna, 2022) which allows us to check not only the continuous parameters of the model but also the tree topology for convergence. Additionally, using a subset of models (Mk, Mk+ACRV *m* = 4, covariomorph *m* = 4) and simulated datasets, we compared the posterior of tree topology differences between the models using `convenience`. In this test, when all the clades compared between two models stay within the threshold for the maximum expected difference in split frequency (caused by sampling randomness), it indicates that the two models explored the same posterior distribution of tree topologies. On the other hand, if some topology splits fall outside the threshold, it suggests that the models explored different posterior of tree topologies.

### 2.4 Empirical Investigation

In our empirical investigation, we examined the performance of our proposed covariomorph model across a diverse range of phylogenetic datasets. We obtained 200 matrices including extant and fossil species from Morphobank (O’Leary and Kaufman, 2012) and Graeme Lloyd’s repository of character matrices (Wright et al., 2016). These matrices are of varying characteristics, including different number of character states (2 states to 8 states), total number of characters (11 characters to 785 characters), and taxonomic sampling (6 taxa to 138 taxa) from various published studies. Our approach was designed to achieve two primary objectives: first, to assess the model’s performance under a diverse collection of datasets, and second, to evaluate the presence of covarion-like rate variation in published studies and its impact in the tree estimation. To accomplish this, we conducted a series of analyses with varying covariomorph model settings.

We examined several sets of rate category configurations, specifically employing 2, 4, 6, 8, 10, and 12 rate categories. With these empirical analyses, we sought to detect potential switching between rate categories; a phenomenon that could signal the presence of covarion-like evolutionary dynamics within empirical phylogenetic matrices. To compare our covariomorph model with the standard models, we analyzed the same matrices using the standard Mk model, both with and without among character rate variation. For the among character rate variation analyses, we set up a discrete lognormal distribution to specify rates maintaining consistency with our covariomorph analyses. We explored two rate category configurations: four (*m* = 4) and eight (*m* = 8) discrete rate categories. In all analyses, the datasets were partitioned by number of states which ensured that the Q-matrices for the data subset containing characters with equal number of states are sized with the observed maximum number of states (Khakurel et al., 2024). For the partitions with appropriately sized covariomorph rate matrices, both the standard deviation and the switching rate is shared across partitions.

The MCMC analyses were run similar to the simulation settings with four independent MCMC runs and for 100,000 generations. At the end of 100,000 MCMC generations, we checked the MCMC runs for convergence using the R package `convenience` (Fabreti and Hö hna, 2022). In the instances where convergence was not achieved within 100,000 generations, we extended the generations to 300,000. If the analysis did not converge even with 300,000 MCMC generations, we discarded that dataset. This resulted in a total of 164 datasets that were analyzed with the selected models mentioned above (Supplementary References; available on Dryad).

#### 2.4.1 Case Studies: Rays and Sharks

##### Datasets

To explore the covariomorph model’s behavior in more detail, we focused on two representative morphological datasets that exhibited rate variation and switching between the rate regimes:

- Rays dataset: Myliobatiformes from Marramà et al. (2023), comprising 52 taxa and 124 characters, and
- Sharks dataset: Neoselachians from Shirai (1996), comprising 53 taxa and 105 characters.

Both datasets are of typical size for morphological datasets, include multistate characters and contain a mix of both extinct and extant taxa. We expanded the number of rate categories for these two datasets, to include 14, 16, and 18 rate categories.

##### Model Selection

Apart from the behavior of the covariomorph model in estimating various model parameters for the rays and sharks datasets, we also assessed the optimal number of rate categories (*m*) for the covariomorph model using marginal likelihood estimation via the stepping-stone (Xie et al., 2011) and path-sampling (Lartillot and Philippe, 2006) algorithms. We compared a series of covariomorph models with varying rate categories (*m* = {2, 4, 6, 8, 10, 12, 14, 16, 18}) along with the standard Mk model and a 4-category among-character rate variation model (Mk+ACRV). We ran power posterior analyses to compute marginal likelihoods using the stepping-stones and path-sampling algorithms (Hö hna et al., 2021). For each analysis, we utilized 128 stepping-stones, discarding an initial 20,000-generation burn-in and subsequently sampling 2,000 generations from each stone. We then computed log Bayes factor (lnBF) for each model against the baseline Mk model.

##### Tree topology comparison

Similar to the simulated datasets, to check whether different models explore different posterior distributions of tree topologies, we utilized `convenience` (Fabreti and Hö hna, 2022) to perform the expected difference in split frequencies test similar to how convergence is assessed. In this test, we compare the expected difference in split frequencies between the posterior samples of trees from different models. If the posterior probabilities for all clades fall within the sampling randomness threshold, we conclude the models explored the same distribution of topologies. Conversely, if clades fall outside this threshold, the models explored different posterior distributions of topologies.

## 3 Results

### 3.1 Simulation Results

Our simulation study demonstrates the efficacy of our covariomorph model in recovering underlying parameters, but crucially reveals that this performance is highly dependent on the total tree length of the phylogeny. As illustrated in Fig. 3, the covariomorph model effectively recovers the true standard deviation (*σ* = 1.174) under the conditions where the data generating model included rate variation for all sets of tree lengths (all conditions except Mk model).

**Figure 3.**
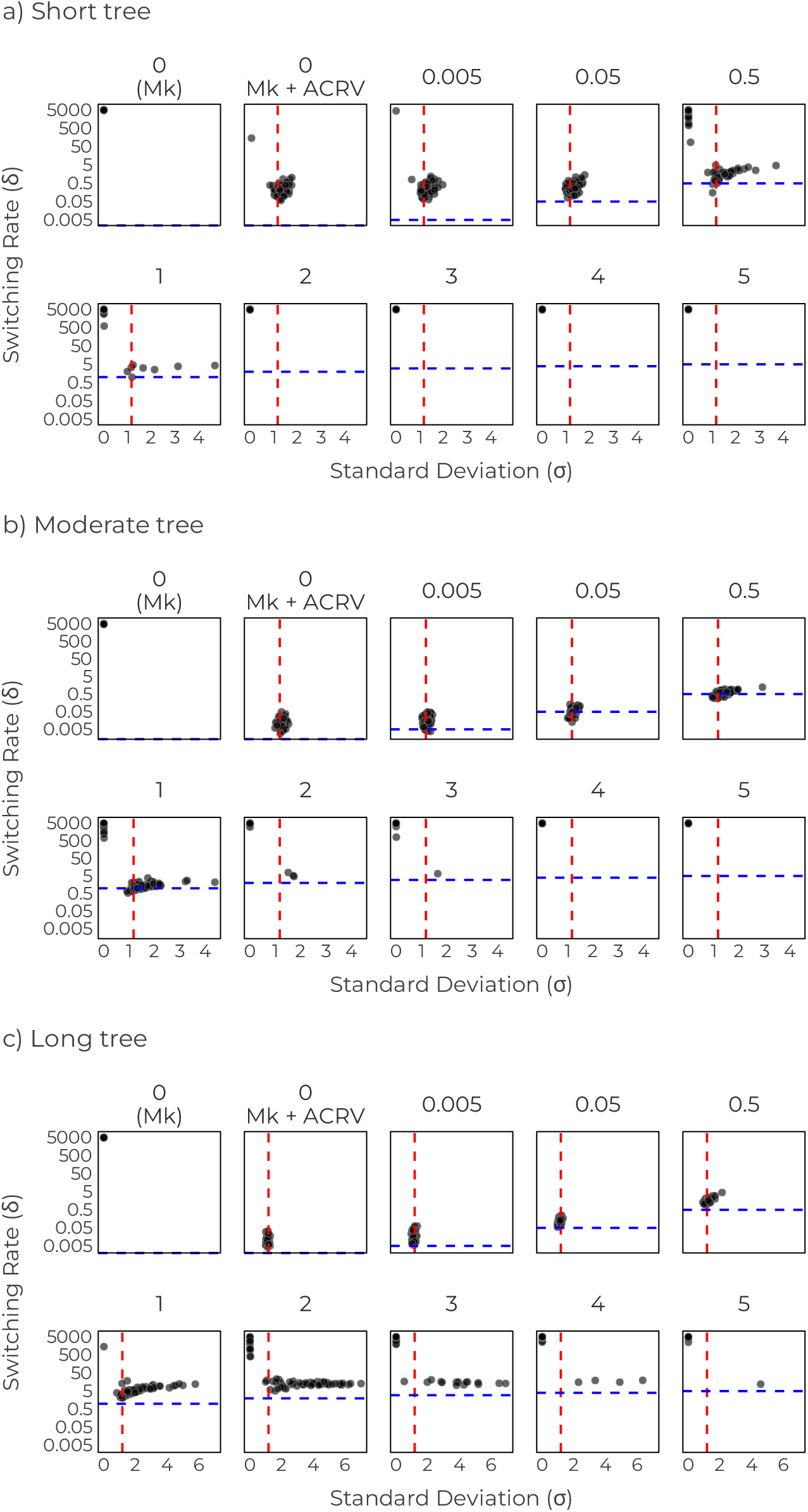
**Posterior median estimates of switching rates (**δ**) and standard deviation (**σ) from 100 simulations under the covariomorph model with four rate categories. Each point represents the posterior median estimate from one simulation replicate. The three rows correspond to simulations on phylogenies with different total tree lengths (TL): (a) Short tree with TL = 3.18. (b) Moderate tree with TL = 10.54. (c) Long tree with TL = 16.71. Each column panel corresponds to a different true switching rate used in simulation, indicated by the panel label at the top. The horizontal dashed lines show the true switching rates for each panel, and the vertical dashed line indicates the true standard deviation used across all simulations (σ = 1.174). The covariomorph model successfully recovers the true parameters across reasonable range of parameters used for simulation depending on the total tree length (see text for more details).

As we expected, for all simulated datasets generated under the Mk model, the covariomorph model defaults to the prior mean for the switching rate *δ*. Additionally, in these datasets, there is no signal of rate variation among characters, thus all the rate categories exhibit equivalent rates (*σ* ≈ 0). This pattern is reflected in the top-left panels of Figures 3a, 3b, and 3c where the simulated replicates converge on a high switching rates near the mean of the prior distribution. Note that for the prior on switching rate, we used a uniform distribution ranging from 0 to 10^4^. This confirms the model’s tendency to collapse to the simplest explanation when no rate heterogeneity is present in the datasets.

In contrast, datasets generated under the among character rate variation model (Mk+ACRV) reveal distinct rate variation between characters without transitioning between different rate categories over time. In these instances, the covariomorph model accurately recovers the true data-generating standard deviation *σ*, reflecting the inherent rate variation present in the data. For the moderate and long trees (Figs. 3b, and 3c; ‘0 (Mk+ACRV)’ panel), the inferred switching rate remains low but non-zero (*δ <* 0.1), consistent with the absence of transitions between the rate categories in the generating process. Longer trees appear to provide slightly more power to constrain the switching rate closer to its true value of zero in this scenario. That is, we inferred *<* 0.1 rate transitions per character state transition, and on average *<* 1 rate transitions across the entire phylogeny (as the tree lengths for moderate and long trees were 10.54, and 16.71 respectively). For the short tree (Fig. 3a, ‘0 (Mk+ACRV)’ panel), the majority of the replicates still indicate a low switching rate (*δ* ≈ 0.22). Even at this rate, the average number of rate switches per character across the tree remains low (approximately 0.8 rate switches on average), confirming the model’s general tendency to correctly constrain the switching rate close to zero when no signal for switching process is present.

For the scenarios involving covariomorph behavior, i.e., where a non-zero switching rate (*δ >* 0) was used to generate datasets, the reliability of parameter recovery depends strongly both on the true switching rate and the total tree length. The model can only reliably distinguish between different rate categories if enough character state changes are observed within each category to estimate its specific rate, compared to how often the process switches between categories. Intuitively, the number of realized transitions required to identify a rate switch depends on the contrast between the distinct rate categories and the duration of time spent in the new regime. If the rate of character state transitions implies stasis (rate ≈ 0), a single or very few realized transitions are sufficient to signal a switch to a non-zero rate category. Conversely, distinguishing between less distinct rates (e.g., ‘medium’ vs. ‘fast’) requires a lineage to persist in the new category long enough to accumulate a cluster of transitions that would be improbable under the original rate. Thus, shorter trees or high switching rates obscure the signal because lineages rarely spend enough time in a single regime to generate the distinct data patterns necessary to diagnose a switch.

Recall that our covariomorph rate matrix was constructed in a way such that the switching rate is in relative units to the expected number of observed character state transitions, thus for a switching rate *δ <* 1.0, more state transitions than rate changes are expected *a priori*. At very low true switching rates (e.g., *δ* = 0.005), few switches occur across the phylogeny, and for all sets of tree lengths, the model’s inference closely resembles that of a model incorporating only among-character rate variation. As the true switching rate increases from these low levels, the precision of the switching rate estimates initially improves.

However, the ability to recover parameters degrades differently depending on the tree length as *δ* increases further. For the short tree (Fig. 3a), the recovery of switching rate parameter (*δ*) degrades quickly as true *δ* increases. When the true switching rate is 0.05, the model recovers both the true *σ* and *δ* robustly. But as the true switching rate increases to 0.5, around half of the simulation replicates fail to recover the true covariomorph behavior, instead collapsing towards Mk-like behavior. In contrast, moderate and long trees (Figs. 3b, and 3c), provide more character state changes overall, allowing the model to tolerate faster switching before the signal averages out. For the moderate tree, reliable recovery extends to *δ* = 1, with degradation becoming pronounced for *δ* ≥ 2. The long tree shows robust recovery up to *δ* = 2, with degradation starting around *δ* = 3.

Thus, while longer trees provide more power to resolve dynamics across a range of switching rates, including higher ones, power decreases for any tree length when the switching rate becomes excessively high relative to the number of character state transitions occurring between switches. In such cases, the process averages across rate categories faster than the signal for rate differences can accumulate. When this occurs, the covariomorph model tends to infer negligible rate variation among characters (*σ* ≈ 0) and high switching rates, effectively collapsing to the behavior expected under the simpler Mk model. This collapse is an expected outcome, as an extremely high switching rate implies many more rate shifts than transitions in the observed character, effectively averaging over the rate categories (which have an average of 1 by design).

For the topology recovery comparison in the simulated datasets, as we expect, when the simplest (Mk) was the true model, the Mk+ACRV and the covariomorph model explored essentially the same tree topologies as the true model (Supplementary Fig. S6a). Specifically, the Mk model failed to recover the topology, and exhibited significantly different clade posterior probabilities than the true generating model, when data were generated under the Mk+ACRV model; likewise, both the Mk and Mk+ACRV models failed when data were generated under the covariomorph model (Supplementary Figs. S6b, S6c). Conversely, the covariomorph model demonstrated flexibility, successfully recovering the correct topology when the true model was with Mk+ACRV or covariomorph, reinforcing its status as a robust generalization capable of accommodating both static and dynamic heterogeneity.

### 3.2 Empirical Results

Our empirical analyses reveal that covarion-like rate variation, where evolutionary rates of characters can change across the phylogeny, is not universal across morphological datasets (Supplementary Fig. S7). The distribution of inferred parameters, specifically the standard deviation of rate variation (*σ*) and the rate switching parameter (*δ*), reveals distinct patterns of character evolution, as visualized in Fig. 4 across models using different numbers of rate categories (*m* = 2, 4, 6, 8, 10, 12).

**Figure 4.**
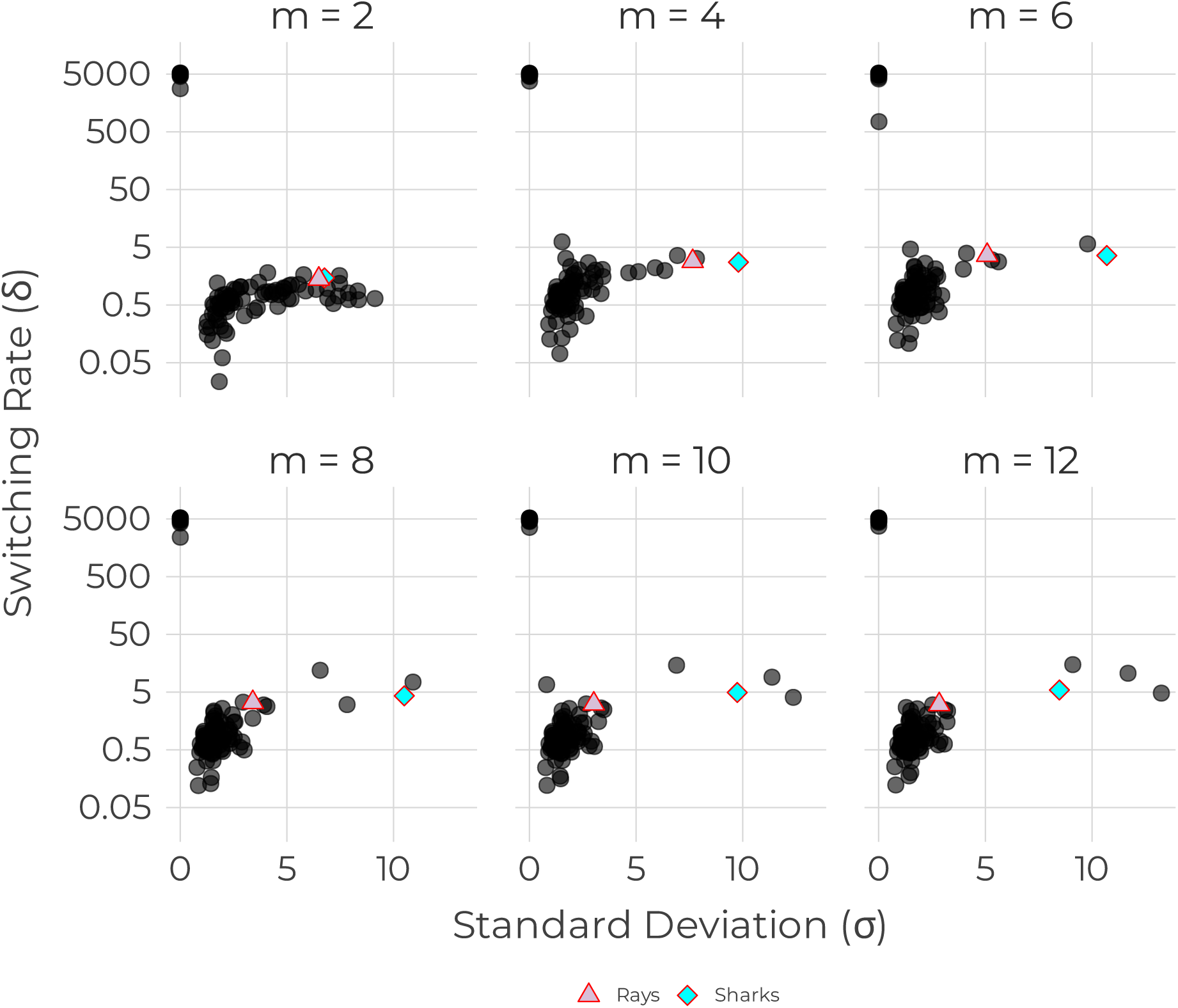
Relationship between inferred switching rates and standard deviation from the covariomorph model with varying rates. Each point represents the median value of the posterior distribution for a specific empirical dataset, aggregated from the analysis of 164 datasets. Two types of empirical datasets can be observed in these plots: first, the ones exhibiting equal rates for all characters (clustered in the top-left); second, showing rate variation between characters (both among character and among lineage). The rays and sharks datasets used for further investigation is indicated by a triangle and diamond respectively.

Fig. 4 shows the posterior median estimates of the switching rate (*δ*, log-scale) against the standard deviation of the lognormal distribution (*σ*) for empirical morphological datasets. Two primary clusters emerge: (i) Approximately half (∼ 77 out of 164) of the examined datasets exhibit parameters clustering near the top-left of the plots (*σ* ≈ 0 and *δ* approaching the prior mean, around 5,000). This indicates negligible rate variation among characters (low *σ*) and extremely high switching between the rate regimes (high *δ*). As shown above in the simulations, such high switching and low standard deviation mimics the behavior of a standard Mk model where all characters evolve under the same rate process. These datasets are more well-explained by equal rates among characters. (ii) The remaining datasets show evidence of rate heterogeneity, characterized by a non-zero standard deviation (*σ >* 0) and a lower switching rate (*δ* generally below 5). These datasets showing rate variation (i.e., *σ >* 0) could exhibit either among-character rate variation or covarion-like dynamics.

The observed distribution underscores the utility of our model. It can effectively accommodate datasets best described by simpler models: reducing to an among-character rate variation model when no switching is detected, or to a standard Mk model when high switching homogenizes rate variation across lineages.

### 3.3 Case Studies: Rays and Sharks

#### 3.3.1 Model selection results

Model selection strongly favors models that account for rate heterogeneity among lineages over the standard Mk model in both rays and sharks datasets (Supplementary Table S1, Supplementary Fig. S8).

For the rays dataset, the Mk+ACRV model showed a negligible improvement over the standard Mk model (lnBF = 1.00). In contrast, all covariomorph models were decisively favored. The simplest covariomorph model (*m* = 2) showed a massive improvement over the Mk model (lnBF = 19.13). Increasing the number of rate categories from *m* = 2 to *m* = 4 resulted in a further substantial improvement in model fit (lnBF increases from 19.13 to 29.41), providing very strong evidence (Kass and Raftery, 1995) for the *m* = 4 model. However, further increase in the number of rate categories (*m* = 6 to *m* = 18) yielded only small changes in the log Bayes factor against the Mk model, which remained relatively stable (between 32.38 and 33.58). This suggests that while modeling rate heterogeneity is important, a model with rate categories higher than *m* = 6 does not significantly improve the marginal likelihood in this dataset.

On the other hand, for the sharks dataset, the preference for models with increased rate heterogeneity was even more pronounced. The Mk+ACRV model was not supported, performing slightly worse than the standard Mk model (lnBF = −1.68). However, the covariomorph models were strongly supported. The log Bayes factor against the Mk model increased sharply from 19.54 (*m* = 2) to 39.88 (*m* = 4) and continued to rise with additional categories, reaching 45.72 for *m* = 6, and appearing to asymptote around a value of 50.27 to 54.23 for *m >* 8. This pattern indicates decisive evidence favoring models with a large number of rate categories *m* ≥ 8 over models with fewer rate categories, suggesting that accurately modeling rate variation in the sharks dataset requires a higher degree of model complexity compared to the rays dataset.

#### 3.3.2 Rate variation and rate category switching

Analysis of the posterior distribution of key parameters of the covariomorph model reveals distinct patterns of rate heterogeneity and model behavior between the rays and sharks datasets, particularly concerning the standard deviation of rate variation (*σ*) and the switching rate (*δ*) as the number of rate categories changes (visualized in Fig. 5).

**Figure 5.**
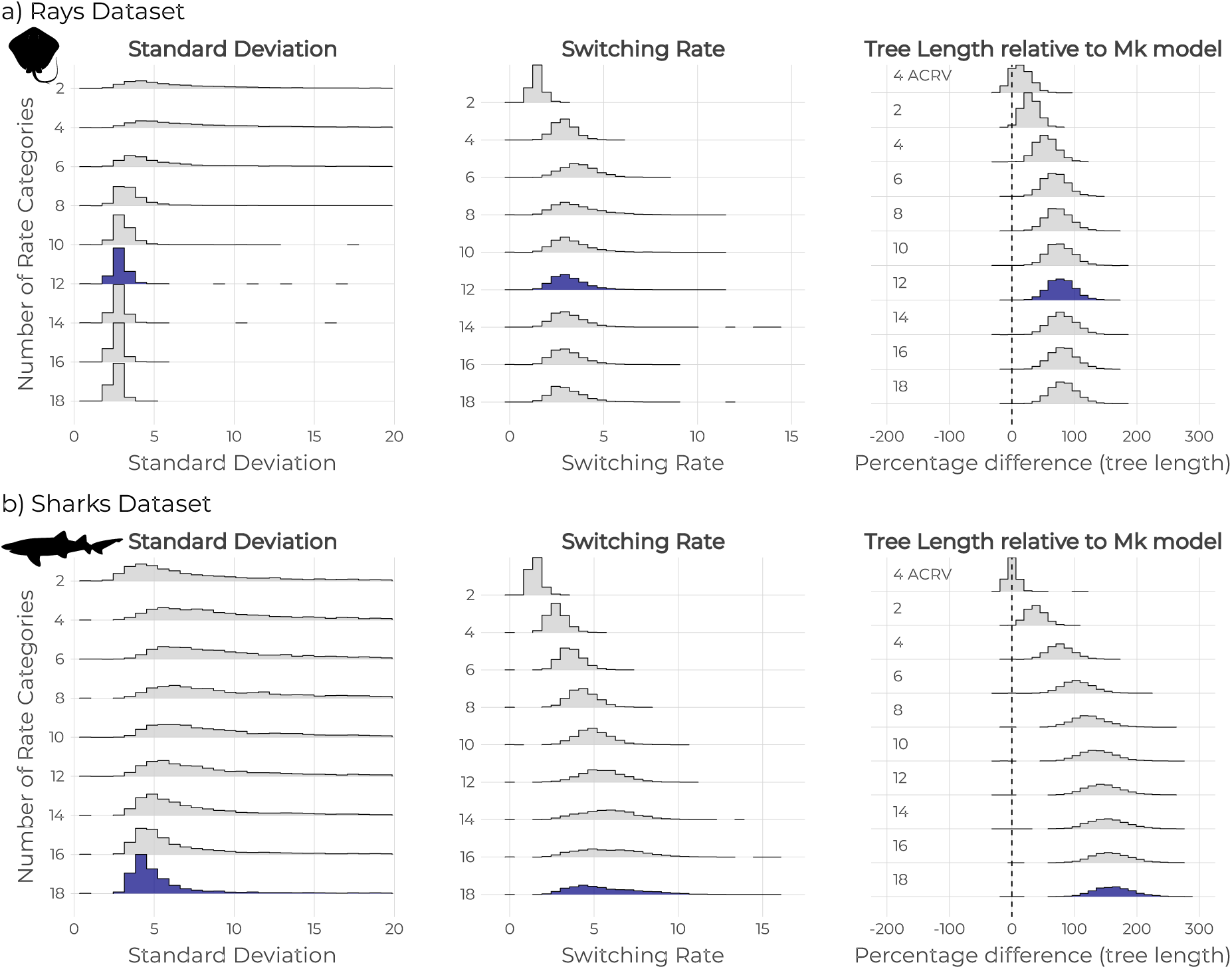
Posterior distributions illustrating the effect of varying number of rate categories on parameter inference for the (a) rays and (b) sharks datasets. From left to right, the panels display the inferred standard deviation, the switching rate between categories, and the percentage difference in tree length relative to the Mk model (indicated by dashed line at 0). Each row represents a separate analysis with different number of rate categories, from 2 to 18. The tree length panel additionally shows the posterior from a 4-category among character rate variation (ACRV) model for reference. The model with highest marginal likelihood is indicated using a darker shade. Silhouettes obtained from PhyloPic (https://www.phylopic.org).

For the rays dataset, posterior estimates for *σ* stabilized as model complexity increased (Fig. 5a, left panel). In models with 10 or more rate categories (*m* ≥ 10), the posterior distributions converged on a stable median *σ* value of approximately 2.8, suggesting that a model of this complexity is sufficient to adequately capture the rate variation present in this dataset. In contrast, the sharks dataset revealed a consistent and robust signal for substantial rate heterogeneity across all model complexities (Fig. 5b, left panel). The median posterior estimate for *σ* initially increased, reaching up to 10.6 for 6 rate categories and gradually decreased as the number of rate categories increased further.

The switching rate (*δ*) shows distinct patterns across the two datasets as the number of rate categories increases (Fig. 5, middle panels). In the rays dataset (Fig. 5a), the posterior distribution of switching rate shows an initial increase with additional rate categories before stabilizing to a slightly lower estimate. The median values of the switching rate ranges from 1.41 for 2-rate category model to 2.97 for the 18-rate category model, reaching up to 3.65 for the 6-rate category model. The posterior uncertainty, represented by the width of the distributions, appears to follow a similar pattern, widening up to 12-rate categories model before narrowing with subsequent addition of rate categories. On the other hand, the sharks dataset (Fig. 5b) exhibits slightly different pattern in the posterior estimates of switching rate with different number of rate categories. As the number of rate categories increased from 2 to 18, the posterior distribution’s location progressively shifts towards higher switching rates, with the posterior median increasing from 1.47 to 5.23. This shift is accompanied by a corresponding increase in posterior uncertainty, as indicated by the widening distributions.

#### 3.3.3 Tree topology and tree length. –

Incorporating different models of rate variation reveal notable impacts not only on the continuous parameters such as the standard deviation or the switching rate but also on the inferred phylogenetic relationships, particularly when accounting for heterotachy. Fig. 6 provides a comparison of the posterior probabilities of splits explored by the Mk model (equal rates), an ACRV model (*m* = 4, character rates vary and are constant across the tree), and the covariomorph model (*m* = 4, rates for characters can vary differently along the branches). For a more comprehensive comparison of tree topology between all the models tested in this study, readers are suggested to view the online supplementary material (Supplementary Figs. S9, S10 and S11).

**Figure 6.**
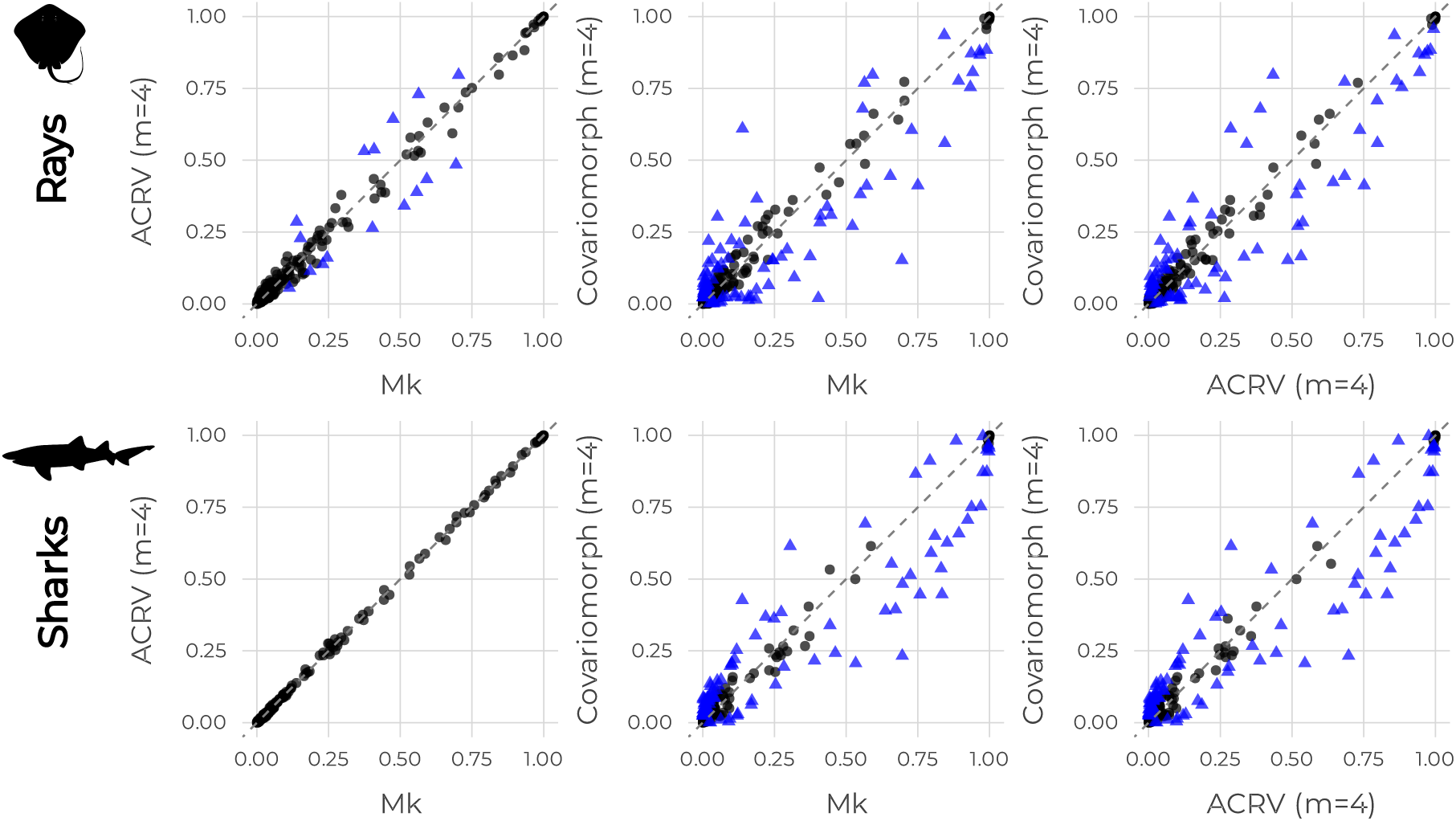
Differences between trees sampled using different models of rate variation for the rays and sharks datasets. These scatter plots show the posterior probabilities of clades, illustrating the level of support each pair of model provides for the same set of potential clades. Here, we include one representative result from each type of rate varying model (Corresponding to Fig. 1), a full comparison of all the models that were tested can be found in the online supplementary material. Each clade is represented by either a circle or a triangle in the plots. Circles indicate clades where posterior probabilities do not significantly differ under the expected difference of split frequencies test for an effective sample size (ESS) of 200, whereas triangles represent clades with significantly different posterior probabilities. In other words, if a clade is a triangle, it was sampled with a significantly different probability between the two compared models.

As shown in Fig. 6, the impact of accounting for among-character rate variation (ACRV) differs between the two datasets. For the sharks dataset, comparison between the Mk and the Mk+ACRV model yields highly congruent results. The posterior probabilities of the splits are almost perfectly correlated (Fig. 6, bottom left), and differences between the splits are minimal (Supplementary Fig. S11), mostly falling well within the expected different for an effective sample size (ESS) of 200. This suggests, adding a model of among character rate variation does not significantly alter the topological inferences compared to the Mk model for the sharks dataset.

For the rays dataset, while the posterior probabilities of clades between the Mk and ACRV models remain highly correlated, there are some clades that are sampled differently by the two models (triangles in Fig. 6, top-left). This indicates that for the rays dataset, accounting for among-character rate variation has some effect on the sampled topology.

In contrast, for both datasets, comparing the models that lack heterotachy (Mk and ACRV) with the covariomorph model (which accounts for heterotachy) reveals substantial differences. The plots comparing the posterior probabilities of clades in the posterior distribution of models accounting for heterotachy and the ones not accounting for heterotachy (Fig. 6, middle and right columns) show that the sampled topology differs significantly. This demonstrates that accounting for heterotachy has a much stronger impact on the sampled topology for both groups rather than accounting for among-character rate variation.

Additionally, the estimated tree lengths also differ substantially between these models. As shown in the rightmost panel of Fig. 5, all covariomorph models (different number of rate categories) yield longer trees compared to the Mk model for both the sharks and rays datasets. For example, in the rays dataset, the posterior median tree length under the covariomorph model with four rates (8.22) is approximately 1.5 times longer than under the Mk model (5.32). The sharks dataset shows a similar pattern, with covariomorph (*m* = 4) tree length 1.7 times longer than Mk tree length.

## 4 Discussion

The evolution of morphological characters is often complex, with rates varying both among characters and along different lineages within a phylogeny (Skinner, 2010; Clarke and Middleton, 2008; Wang and Lloyd, 2016; Lloyd et al., 2012). Standard models such as the Mk model fail to capture this heterogeneity. In this study, we introduced and evaluated the *covariomorph* model —an extension of the covarion model to morphological data— implemented in `RevBayes`, designed specifically to incorporate character-specific rate variation in discrete morphological datasets.

Our simulations demonstrate that the model can accurately recover underlying parameters, including the rate of switching between evolutionary rate categories (*δ*) and the magnitude of rate variation among categories (*σ*). While we focused our empirical validation on datasets that exhibited complex rate dynamics, we note that the covariomorph framework nests the standard morphological models. As demonstrated in our simulations (Fig. 3), when the switching rate (*δ*) approaches zero, the model effectively collapses to the Mk+ACRV process; similarly, it collapses toward the Mk model when rate variation is absent. Consequently, our simulation study serves to delineate the specific conditions under which the covariomorph model offers a distinct improvement over static alternatives, illustrating that the model behaves appropriately even when the generating process is simpler than the inference model.

Additionally, application to a diverse set of empirical datasets revealed two distinct clusters of model behavior: approximately half exhibited parameters consistent with the simple Mk model, while the remaining datasets showed clear evidence of rate heterogeneity (*σ >* 0) that is compatible with either static among-character rate variation (Mk+ACRV) or dynamic covarion-like processes (covariomorph). Crucially, while our Bayes factor analysis provided decisive evidence for the complex covariomorph model in the rays and sharks datasets (Supplementary Table S1, Supplementary Fig. S8), for the majority of the remaining datasets, the parameter estimates are ambiguous and should not be over-interpreted as definitive proof of covarion dynamics. Instead these ambiguous results indicate that covarion processes cannot be ruled out and offers a plausible alternative explanation for the observed heterogeneity, particularly in datasets with sufficient evolutionary depth where the dynamics are theoretically resolvable. Importantly, incorporating these dynamics significantly impacts the phylogenetic tree inferences, influencing both the tree topology (Fig. 6) and the branch length estimates (Fig. 5) compared to traditional models. This finding has critical implications for downstream analyses that rely on branch lengths, such as divergence time estimation and the calculation of evolutionary rates.

### 4.1 Incorporating conditioning on variable sites

While our study focused on inferring phylogenies from morphological characters, we did not explicitly account for a bias inherent in typical morphological datasets. Morphological datasets are typically curated to include only the traits that differ among the included taxa. This bias, also referred to as “ascertainment bias,” can lead to overestimation of character transition rates and thus branch length inflation (Lewis, 2001). Lewis (2001) proposed a correction for the Mk model, termed the Mkv model, which conditions the likelihood calculation on the observed variability of a character.

Even though ascertainment bias correction is very important for morphological phylogenetics (Mulvey et al., 2024; Capobianco and Hö hna, 2025), we excluded it from our study. Our implementation of the covariomorph model does not currently incorporate this ascertainment bias correction. The main reason is that standard implementations of ascertainment bias correction do not work off the shelf for the covariomorph model. This is because the covariomorph uses character expansion to model the hidden rate categories. Future development should prioritize integrating an ascertainment bias correction into the covariomorph framework, analogous to the mMkv model (Capobianco and Hö hna, 2025). For such an implementation, we suggest that the conditioning on variability should be based on the original observed character states rather than the expanded virtual states used internally by the covariomorph model. Incorporating this correction is a critical next step for real-world datasets and would provide more accurate estimates of evolutionary rates and branch lengths.

### 4.2 Interaction between switching rates and character state transition rates

In the current formulation of the covariomorph model, both the rate of character state transitions (e.g., under the Mk model) the switching rate between the rate categories (*δ*) are governed by a shared *evolutionary clock*. This coupling implies that branches with faster evolutionary rates (i.e., longer branches) are expected to exhibit proportionally more state transitions and more frequent switches between rate categories.

While this formulation simplifies inference and reduces the number of free parameters, it may obscure biologically meaningful differences between the processes of observable character change and hidden rate regime shifts. Rapid morphological evolution in a lineage (e.g., due to ecological shifts or developmental plasticity) does not necessarily imply an increased tendency to shift between distinct evolutionary rate regimes. In other words, fast observable evolution does not necessarily entail high heterotachy. The model’s assumption of proportionality between transitions and switching thus conflates two potentially distinct processes.

Empirical support for this concern is evident in Fig. 5: as the number of rate categories increases, the posterior median of the switching rate (*δ*) often shows a modest but consistent decline, while the estimates of the magnitude of rate variation, i.e., the standard deviation (*δ*) remain relatively stable. This pattern suggests that the model might be compensating for increased complexity (i.e., more rate categories) by suppressing switching, even when observable rate variation persists. Particularly, in the sharks dataset, a stable high *σ* is paired with decreasing *δ* with additional rate categories.

These patterns highlight the need for a model extension that explicitly decouples the character transition rates from rate switching dynamics; potentially by introducing branch-specific switching process or separating scaling parameters. Such an approach would allow the covariomorph model to more accurately represent evolutionary scenarios where the tempo of morphological change is independent from the frequency of heterotachy.

### 4.3 Relaxing Symmetry in Rate Regime Transitions

In the covariomorph model, heterotachy is modeled via hidden rate categories that characters transition among through time, governed by a continuous-time Markov process. Typically, the switching process assumes equal stationary probabilities for each rate category, implying that, over evolutionary timescales, lineages are equally likely to reside in any given evolutionary rate regime.

Relaxing this assumption to allow for unequal category probabilities, that is, differing probabilities *π_i_* for a character being in category *i*, permits the model to capture biologically realistic asymmetries in how often different evolutionary rate regimes are occupied. These unequal probabilities can be obtained either from explicit parameterization (non uniform *π*) or indirectly via asymmetric switching rates (*δ_ij_* = *δ_ji_*).

This is particularly relevant in scenarios where heterotachy is itself structured: for example, transitions from a conserved (slow-evolving) state to an accelerated evolutionary regime may be more common than the reverse. In adaptive radiations, such as that of African cichlid fishes, characters may have entered the high-rate categories frequently —corresponding to bursts of morphological evolution— while reversals to low-rate regimes were rare during the diversification phase (Kocher, 2004; Salzburger, 2018)

Conversely, after key innovations like the evolution of the turtle shell, lineages may have transitioned into low-rate categories and remained there for extended periods, as constraints favored stasis (Lyson et al., 2014). This would imply a higher probability for the slow-rate category, even if the initial transition from a fast rate occurred. Similar dynamics may explain long periods of morphological stasis in lineages like coelacanths (Turner, 2019), where heterotachy is asymmetric; rates slowed and remained low, but rarely re-accelerated.

Incorporating unequal probabilities for different rate regimes thus allows researchers to model not just when rate shifts occur, but how heterotachy is distributed across the phylogeny. This adds biological realism by decoupling the tempo of switching from the frequency of different rate regimes, enabling finer-grained inferences about evolutionary dynamics across deep-time.

### 4.4 Partitioning Characters by Evolutionary Model

In phylogenetics, it is common that the Markov model characterized by an instantaneous rate matrix Q is constant during the entire evolutionary process. While models that relax this assumptions —for instance, by allowing rates to vary among characters (Γ) or across lineages (relaxed clocks)—add realism, they still apply a single type of evolutionary process to all characters in a dataset (Gascuel and Guindon, 2007; Baele et al., 2021). Our work here introduces the covariomorph model as a powerful tool for capturing lineagespecific rate heterogeneity, but our own results suggest that even this flexible model may not be the optimal choice for every character. While the covariomorph model, as implemented here, provides estimates of global parameters that represent the average rate-switching behavior integrated across all characters, it does not provide per-character estimates, and thus cannot distinguish between characters that follow a simple Mk process and those that are highly heterotachous.

A key finding from our empirical investigation of 164 datasets is that they fall into two distinct camps: approximately half showed strong evidence of both among-character and covarion-like rate variation, while the other half were best explained by a simpler, more uniform process, akin to the standard Mk model. This dichotomy at the level of entire datasets strongly implies that a similar heterogeneity of process likely exists within individual datasets. It is biologically plausible that some characters (e.g., highly constrained, functional traits) evolve under a simple, time-homogeneous process, while others (e.g., traits under shifting selective pressures) are better described by a more nuanced covariomorph model.

Therefore, a logical and powerful extension would be to partition the data matrix by the evolutionary model itself. Such an approach would apply the standard Mk model to one subset of characters and the covariomorph model to another. This could be implemented in several ways. An *a priori* approach might involve assigning characters to partitions based on external criteria, such as the belief of the researcher. However, we note that such partitioning schemes are not always straightforward.

A more elegant and data-driven solution would be to use a mixture model framework. This would allow the analysis to infer which character belong to the “simple” (Mk) class and which belong to the “complex” (covariomorph) class, potentially even estimating the proportion of characters in each. By applying the right level of complexity to the right characters, such a mixed-model approach could provide a more accurate and nuanced picture of morphological evolution, likely improving the overall fit and accuracy of the resulting phylogenetic inference.

## 5 Conclusion

In this paper, we implement and test a phylogenetic model that allows for heterotachy in morphological datasets. Our covariomorph model provides a more nuanced framework for analyzing discrete morphological data and potentially reshaping the inference of phylogenetic trees using discrete morphological characters. By explicitly modeling heterotachy —the variation of evolutionary rates for individual characters across different lineages— it offers a more flexible and potentially more realistic representation of morphological evolution by accounting for temporal variation in fitness than standard models assuming rate homogeneity. Our analyses show that while definitive evidence favoring covariomorph dynamics is only present in a few examined datasets, rate variation compatible with a covarion-like process is frequently observed in empirical datasets. Accounting for this possibility is important as it significantly impacts both the tree topology and the branch length estimation, with potential consequences for understanding evolutionary timescales and rates. While further extensions to relax the model assumptions are possible, the covariomorph approach represents a significant step towards capturing the complex, dynamic nature of morphological character evolution.

## 6 Supplementary Material

Supplementary files (supplementary figures and tables) is available at https://github.com/basantakhakurel/morpheus.

## Acknowledgments

We would like to thank the Hö hna Lab at LMU Munich for helpful feedback on an earlier version of the manuscript.

