## Supplementary File for "A covarion model for phylogenetic estimation using discrete morphological datasets"

This document is a supplement accompanying the manuscript ‘A covarion model for phylogenetic estimation using discrete morphological datasets’. It consists of the data sources used in the main manuscript for testing the *Covariomorph* model. In addition to that this document also includes a few introductory figures about the model (Figs. S1, S3, S5, and S4), coverage plots validating the implementation in `RevBayes` (Fig. S2), and some results (Figs. S7, S9, S8, S10, S11).

#### Graphical representation of the Covariomorph model

We can visualize the Covariomorph model by following a graphical model style (Jordan, 2004) introduced in phylogenetics by Höhna et al. (2014). Figure S1 illustrates the graphical model representation of the covariomorph model used in this study. The phylogenetic tree ($\psi$) is defined by a uniformly distributed topology $\tau$ over $N$ taxa and branch lengths $bl_i \sim$ Exponential($\lambda$), where $i \in 2N - 3$ and $\lambda$ is 10. Morphological character evolution is modeled using a phylogenetic continuous-time Markov chain (PhyloCTMC) parameterized by $\psi$ and a covariomorph rate matrix $Q_{cov}$. The covariomorph model introduces heterogeneity in character transition rates via a set of $j$ rate categories, each associated with a rate multiplier $m_j \sim$ LogNormal($\mu_m = 0, \sigma_m$), where $\sigma$ is a hyperparameter controlling the magnitude of rate variation. Switching between the rate categories occur at a uniform rate ($\delta \sim$ Uniform), and within each rate categories, character transitions follow a standard Mk model ($Q_{Mk}$). The combined process $Q_{cov}$ captures both among-character and temporal rate heterogeneity by integrating over the hidden rate process, and is used to generate morphological data or infer the phylogeny from the morphological data $seq$.

#### Validation of Implementation

Validating the implementation of the covariomorph model in `RevBayes` was done by Simulation Based Calibration (Talts et al., 2018). According to this approach, if we draw parameter values from their prior distribution and subsequently simulate data using these parameter values, the inferred credible intervals will contain the true parameter value in the same percent of the repetitions. In our test, we used a ‘true’

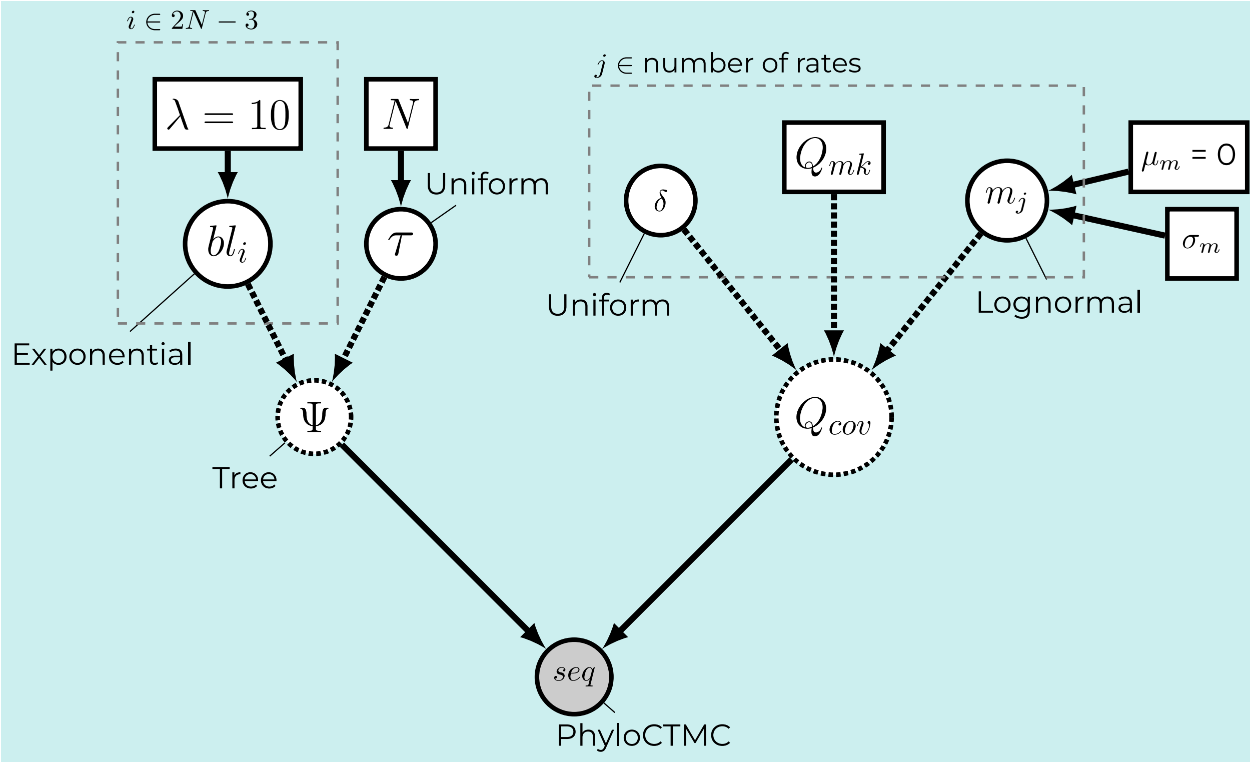


Figure S1: A graphical model displaying the components of the covariomorph model setup in the study. Following Höhna et al. (2014).

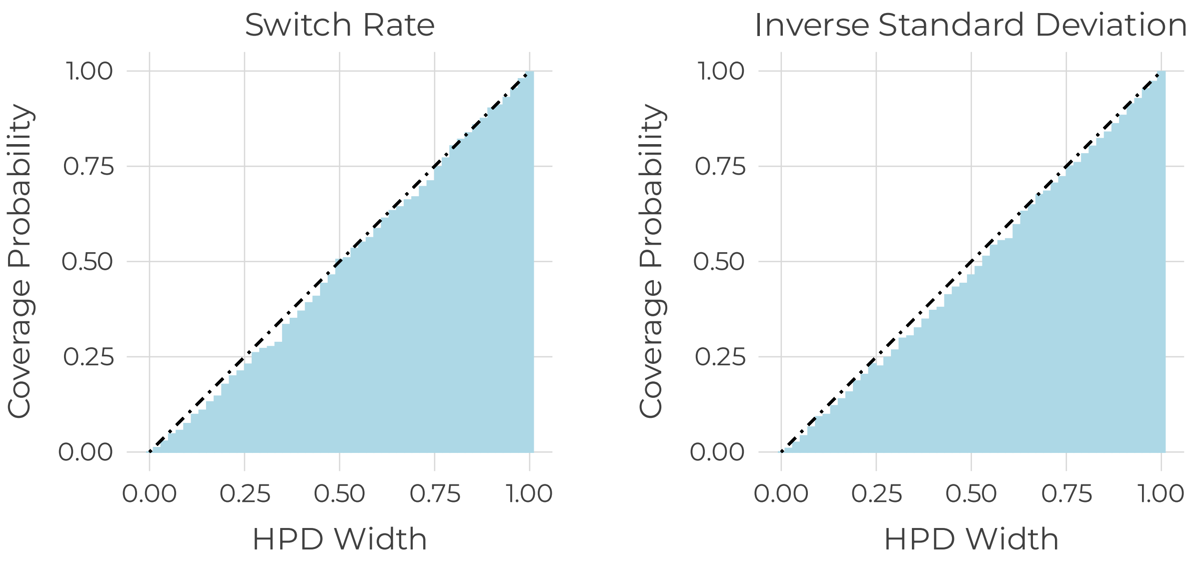


Figure S2: Results of the validation analysis for the covarion model implementation. These plots show how well the highest posterior density (HPD) captures the true parameter values across different interval widths.

value for the inverse of standard deviation of the lognormal distribution and a 'true' value for the switching rate to generate datasets. Upon performing MCMC, the estimated value for the standard deviation should fall within the 95% HPD (highest posterior density) at least 90% of the time for the coverage to be 0.9. The results from our validation analyses is visualized in Figure S2.

#### Virtual states expansion

In the Covariomorph model, the observed character states at the tips of the phylogeny are expanded into "virtual states" to account for different evolutionary rate regimes. Figure S3 illustrates this process for a character with two observed states (0 and 1) and three rate categories. Each observed state is expanded into a set of virtual states, where each virtual state corresponds to the observed state evolving under a specific rate category. For instance, an observed state of '1' is expanded into the set of virtual states $\{1, 3, 5\}$, and an observed state of '0' becomes $\{0, 2, 4\}$. This expansion allows the model to simultaneously infer the character state and the evolutionary rate category at any point in the tree, capturing the heterogeneity of the evolutionary process.

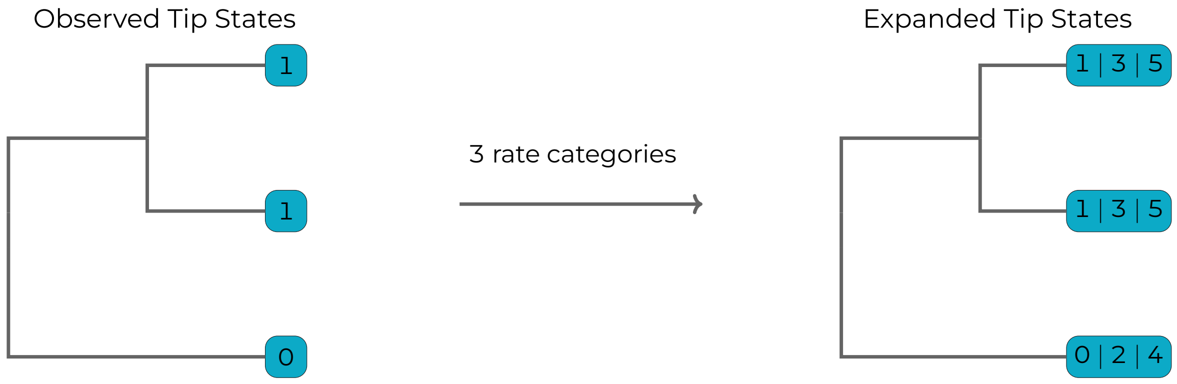


Figure S3: Virtual states obtained from the observed states when the number of rate category is 3.

#### Computational Requirements

The covariomorph model necessarily introduces additional computational costs compared to the standard Mk model. This increase is driven by two factors:

1. **Additional Parameters:** The model adds two new global parameters to be estimated: the shape parameter of the rate distribution ($\sigma$ in our case) and the switching rate ($\delta$). This is one more parameter than the Mk+ACRV model.

2. **Expanded State Space:** The primary increase in computational load comes from the expansion of the instantaneous rate matrix (Q-matrix). The model uses $m$ hidden rate categories, expanding the $k$-state character matrix to a $k \times m$ "virtual state" matrix. All likelihood calculations are performed on this larger matrix.

##### Runtime Analysis

We performed a benchmark analysis to quantify the relative runtime of the covariomorph model compared to the Mk and Mk+ACRV models.

**Methods:**

- We simulated datasets varying in both the number of taxa (10, 20, 30) and the number of characters (50, 100, 200, 300, 400).
- We compared the performance of five model configurations: Mk, Mk+ACRV ($m = 4$), Mk+ACRV ($m = 8$), Covariomorph ($m = 4$), and Covariomorph ($m = 8$).
- All benchmark analyses were performed on a single compute node CPU (AMP EPYC 9654 2.4 Ghz) using `RevBayes` compiled without parallelization.
- Each run consisted of a single MCMC chain for 50,000 generations.

**Results:**

The results of the benchmark are presented in Figure S4.

- The covariomorph model is the most computationally demanding, and its runtime scales significantly with the number of rate categories ($m$).
- With 8 rate categories, the covariomorph model can be over two orders of magnitude slower than the baseline Mk model, particularly on datasets with fewer taxa where the relative cost of the complex Q-matrix calculation is the highest.
- The Mk+ACRV model shows a similar, though less pronounced, increase in cost as $m$ increases.
- While the covariomorph model is computationally intensive, it remains a practical and feasible option for the dataset sizes typically used in morphological phylogenetics.

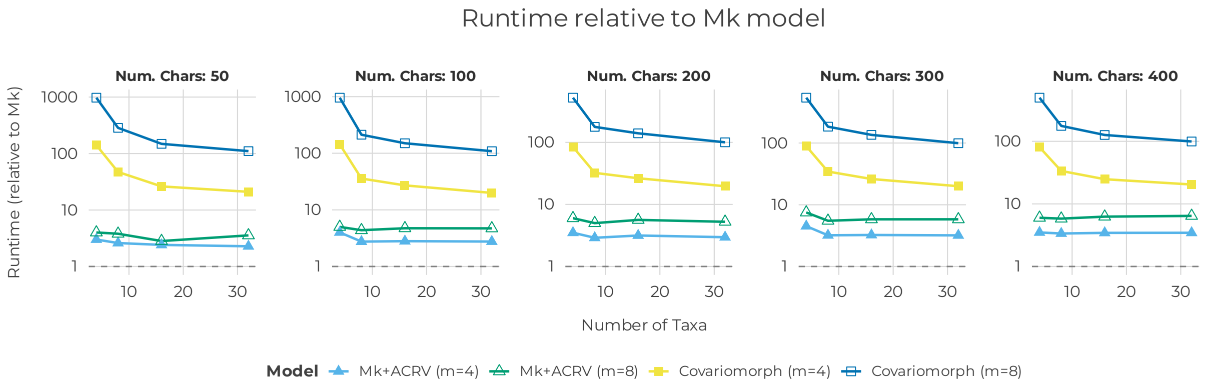


Figure S4: Relative runtime of Mk, Mk+ACRV and Covariomorph models across datasets varying in the number of taxa and characters compared to the baseline Mk model. Values greater than one indicate slower performance relative to Mk. The covariomorph model is consistently more demanding than Mk or Mk+ACRV, especially with higher number of rate categories, but remains practical for datasets of typical size.

#### Memory Requirements

The memory usage for likelihood calculation also scales with the model's complexity. Following Smith et al. (2024), the memory required grows linearly with the number of taxa, the number of site patterns, the number of character states ($k$), and crucially for this model, the number of rate categories ($m$).

This linear scaling is a direct result of the expanded $k \times m$ virtual state space used for the likelihood calculations. For all the datasets analyzed in this study, the memory demands remained well within the limits of a typical modern workstation.

### Discretization of the Lognormal distribution

The covariomorph model utilizes a discretized lognormal distribution to model variation in evolutionary rates across different branches and characters. Figure S5 provides a visual representation of how the rate categories are derived from this distribution. This figure illustrates the relationship between the number of discrete rate categories $k$ and the shape parameter $\sigma$ of the lognormal distribution. The plots demonstrate two key points:

- As the shape parameter $\sigma$ (x-axis) increases, the variance among the rate categories also increases, leading to a wider spread between the fastest and slowest rates.
- As the number of rate categories $k$ (different facets) increases, the continuous lognormal distribution is approximated more finely.

But it comes with a caveat. A notable consequence of using may rate categories (e.g., $k \geq 10$) is that a substantial portion of these categories are assigned vary low rate values ($\leq 0.1$). Since each category is equiprobable, this can lead to an unintentional up-weighting of slower rate categories.

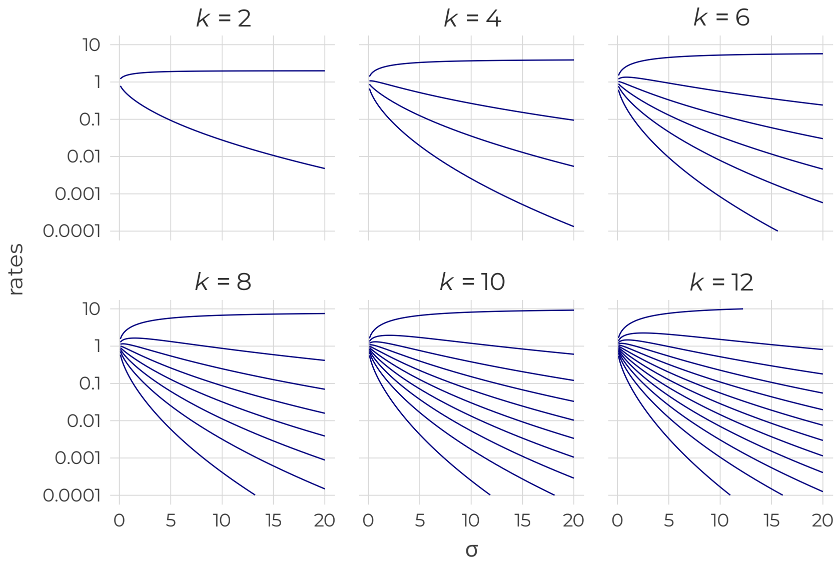


Figure S5: Rate categories from a discretized lognormal distribution. Each panel shows the values of the discrete rate categories as a function of the distribution's shape parameter, $\sigma$, for a different number of total categories ($k$). Within each model, the category rates (y-axis) are chosen to have equal probability ($p = 1/k$).

#### Topological Difference in Simulated Datasets

To evaluate the impact of model misspecification on tree topology, we conducted comparison test on simulated datasets. Using `convenience` (Fabreti and Höhna, 2022), we performed the expected difference in split frequencies test similar to how convergence is assessed. We specifically chose dataset generated under three distinct true models:

1. Mk model ($\sigma = 0,\ \delta = 0$): Data generated with simple time-homogeneous rates.
2. Mk + ACRV model ($\sigma = 1.174,\ \delta = 0$): Data generated with static rate variation across characters.
3. Covariomorph model ($\sigma = 1.174,\ \delta = 0.5$): Data generated with dynamic rate variation and switching between the rate categories.

Each of these datasets were independently analyzed under all three inference models (Mk, MK+ACRV, and Covariomorph) using the same MCMC settings described in the main manuscript.

We assessed the similarity of the posterior tree distribution inferred by each model using the expected difference in split frequencies test. This test determines if the posterior probability calculated for a given clade (split) differs significantly between two models, beyond what is expected from Monte Carlo sampling error. Clades falling outside the threshold (defined by an Effective Sample Size, $ESS = 200$ in our test) are considered to have significantly different posterior probabilities, indicating that the two models explored different regions of the tree topology space.

Figure S6 demonstrates the posterior probability of each unique clade observed in the joint posterior distribution for two compared models. Clades with significantly different posterior probabilities are highlighted, providing a direct visual representation of topological discordance under model misspecification.

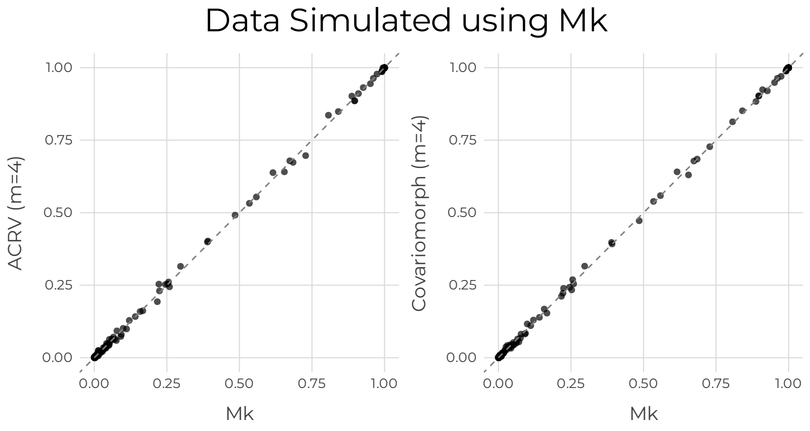


(a) Clade posterior probabilities when the true model is Mk

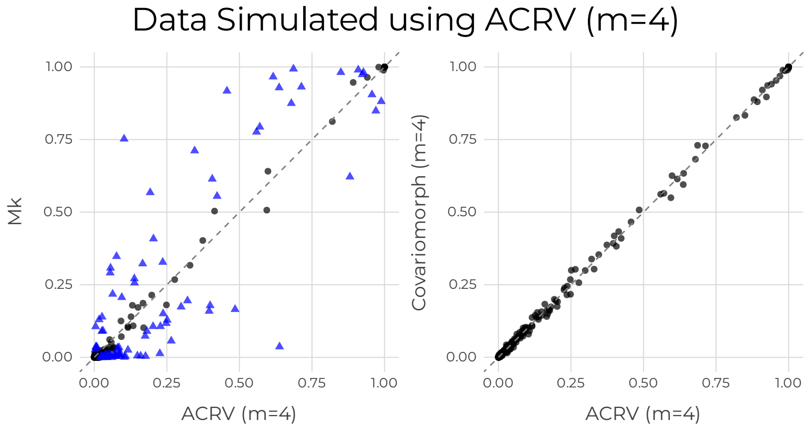


(b) Clade posterior probabilities when the true model is Mk + ACRV ($m = 4$)

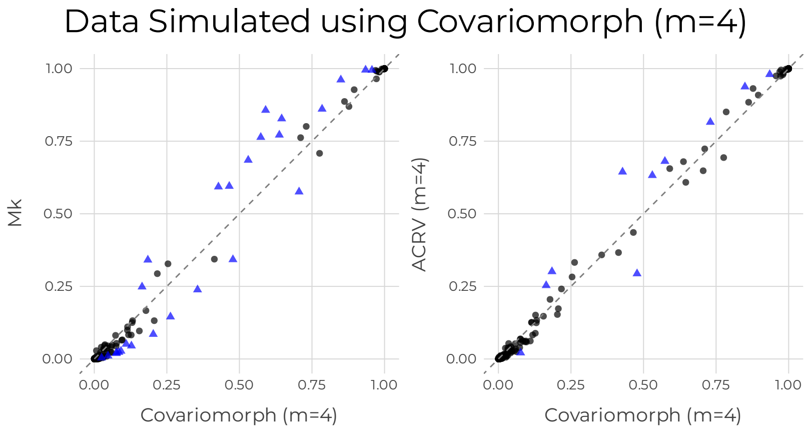


(c) Clade posterior probabilities when the true model is Covariomorph ($m = 4$)

Figure S6: Clade posterior probabilities for different true models using simulated datasets.

#### Empirical datasets: Size and parameter estimates

Figure S7 displays the landscape of the empirical morphological datasets used in this study, plotting the number of taxa against the number of characters for each dataset. The datasets vary considerably in size, ranging from just a few taxa and characters to over 100 taxa and hundreds of characters.

The plots are colored according to the values of key parameters inferred by the covariomorph model with four rate categories:

- Standard Deviation, $\sigma$ (Panel a): This plot shows the inferred standard deviation of the lognormal distribution that indicates the magnitude of rate heterogeneity. There is a slight correlation between the dataset size and the inferred value of $\sigma$, with low values appearing mostly for datasets with lower number of taxa and characters.

- Switching Rate, $\delta$ (Panel b): This plot illustrates the inferred rate of switching between different evolutionary rate categories. A potential trend can be observed where datasets with fewer characters and taxa (yellow points) are better explained by simpler model, such as Mk. This suggests that a larger number of characters may be necessary to provide sufficient information for the model to infer meaningful rate switching across the phylogeny.

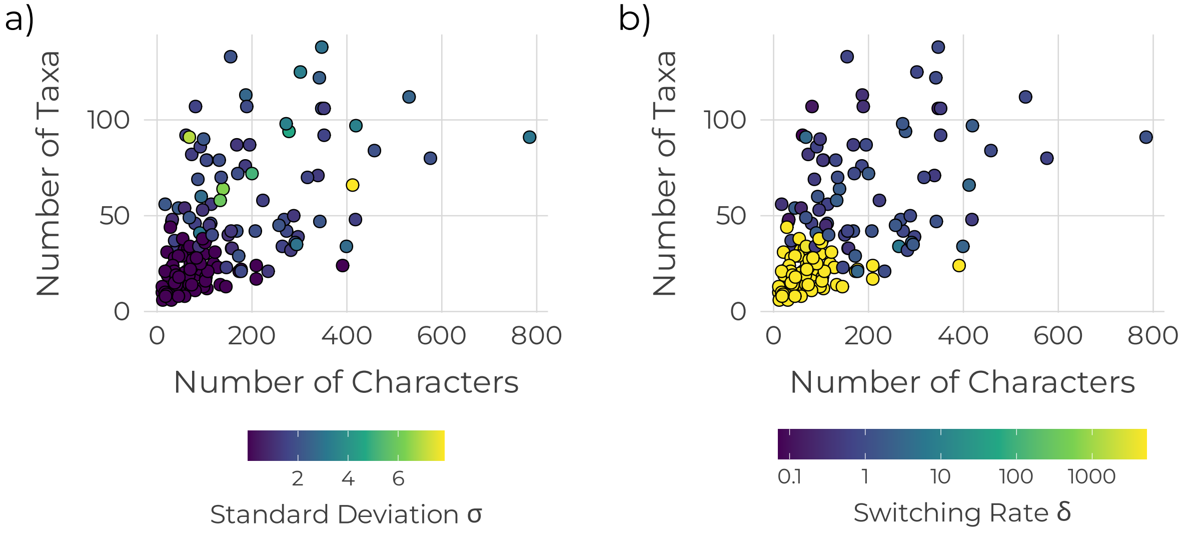


Figure S7: Plots showing the size of morphological dataset used in this study. Each point represents a dataset with corresponding number of characters and taxa colored by a) Standard deviation ($\sigma$) and b) Switching rate ($\delta$) inferred from Covariomorph model with 4 rate categories.

#### Comparison between the Models

To evaluate the performance of the Covariomorph model, we compared it against the standard Mk model and the Mk model with among-character rate variation (Mk+ACRV). The comparison was conducted using model selection based on marginal likelihood estimates. For the covariomorph model, we explored a range of different numbers of discrete rate categories ($m$), from 2 to 18, to assess how model fit is influenced by the number of rate categories allowed.

The marginal likelihoods for each model and for each number of rate categories were estimated for the Rays (Marramà et al., 2023) and Sharks (Shirai, 1996) datasets, as shown in Figure S8 and Table S1. Bayes factors ($ln(BF)$) were calculated to compare the evidence for each model against the baseline Mk model. As seen in the results, the covariomorph model consistently provides a better fit to the data than both the Mk and Mk+ACRV models for both datasets, with the marginal likelihood generally increasing as more rate categories are added, before reaching a plateau. This indicates strong support for a model that allows for rates to vary across both characters and lineages.

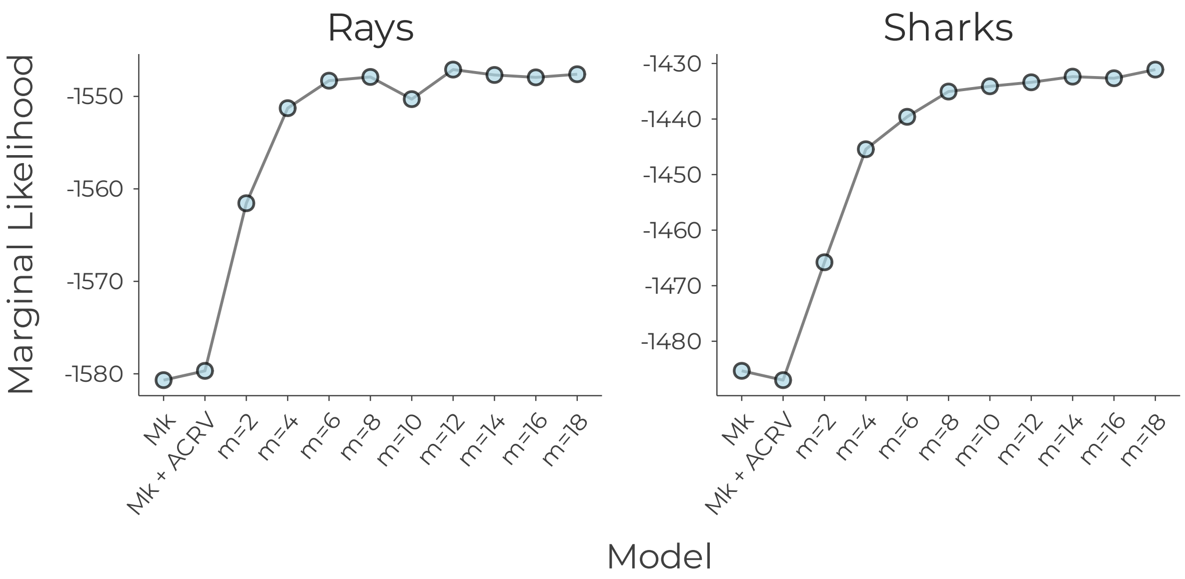


Figure S8: Marginal Likelihood estimates of Covariomorph model with varying number of rate categories

Table S1: Results of model selection analysis on Rays and Sharks dataset with varying number of covariomorph rate categories $m = \{2, 4, 6, 8, 10, 12, 14, 16, 18\}$. Bayes factor $ln(BF)$ is calculated against the Mk model.

| Dataset | Mk model $\ln(L)$ | Bayes factors $ln(BF)$ for number of rate categories ($m$) | | | | | | | | | |
|---|---|---|---|---|---|---|---|---|---|---|---|
| | | Mk+ACRV ($m = 4$) | 2 | 4 | 6 | 8 | 10 | 12 | 14 | 16 | 18 |
| Rays | -1580.68 | 1.00 | 19.13 | 29.41 | 32.38 | 32.79 | 30.39 | 33.58 | 33.00 | 32.74 | 33.08 |
| Sharks | -1485.32 | -1.68 | 19.54 | 39.88 | 45.72 | 50.27 | 51.22 | 51.94 | 52.96 | 52.67 | 54.23 |

#### Topological difference

To evaluate the topological difference between the trees inferred by the different model, we calculated the normalized Robinson-Foulds (RF) score (Robinson and Foulds, 1981) between the maximum *a posteriori* tree (MAP) (Heled and Bouckaert, 2013) from each model and the MAP tree from the Mk model for both the Rays and Sharks datasets. The RF score measures the dissimilarity between two phylogenetic trees, where a lower score indicates a higher degree of similarity.

The results in Figure S9 reveal how the choice of the model affects the topology of the inferred tree relative to the standard Mk model.

- For the Rays dataset, the RF scores were generally higher for all models, indicating a greater overall dissimilarity from the Mk model tree. Here, the Mk+ACRV model's tree showed the least dissimilarity compared to the covariomorph models.
- For the Sharks dataset, a different pattern emerged, the tree inferred using the Mk+ACRV model showed the least difference from the reference phylogeny, with the lowest RF score. The trees from the covariomorph models all exhibited greater topological distances from the Mk tree, with RF scores ranging from approximately $0.28$ to $0.40$.

Additionally, Figures S10 and S11 show the topological difference across the entire posterior sampled from different models. A clear trend that can be observed is that, the models without heterotachy (Mk and Mk+ACRV) are congruent with each other and the covariomorph models (with varying number of rate categories) are congruent with each other. On the other hand, the models without heterotachy and the covariomorph model are exploring different tree topologies based on the expected difference in split frequencies with a threshold of $ESS = 200$.

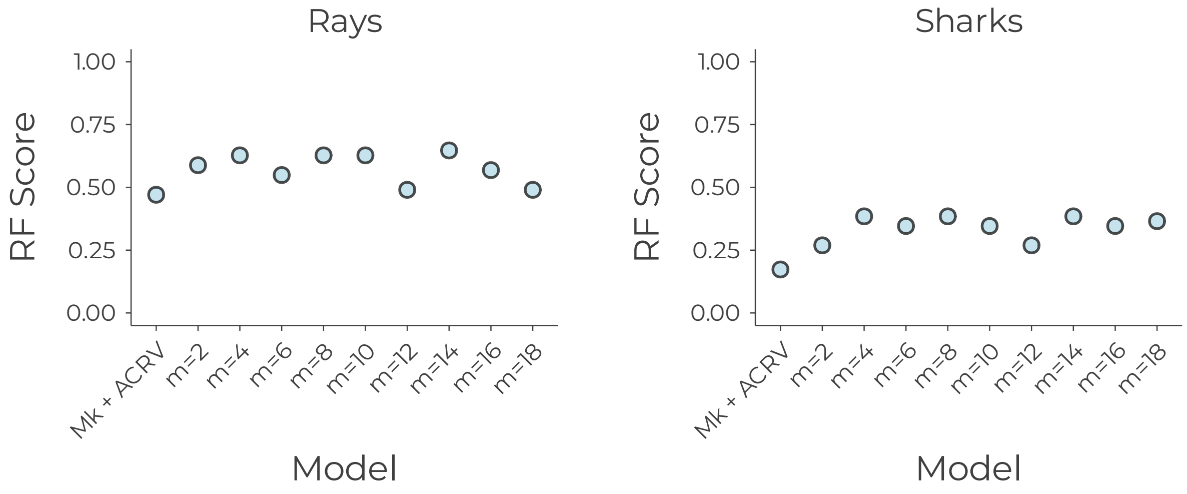


Figure S9: Normalized Robinson Foulds (RF) scores between the MAP trees for the rays and sharks datasets.

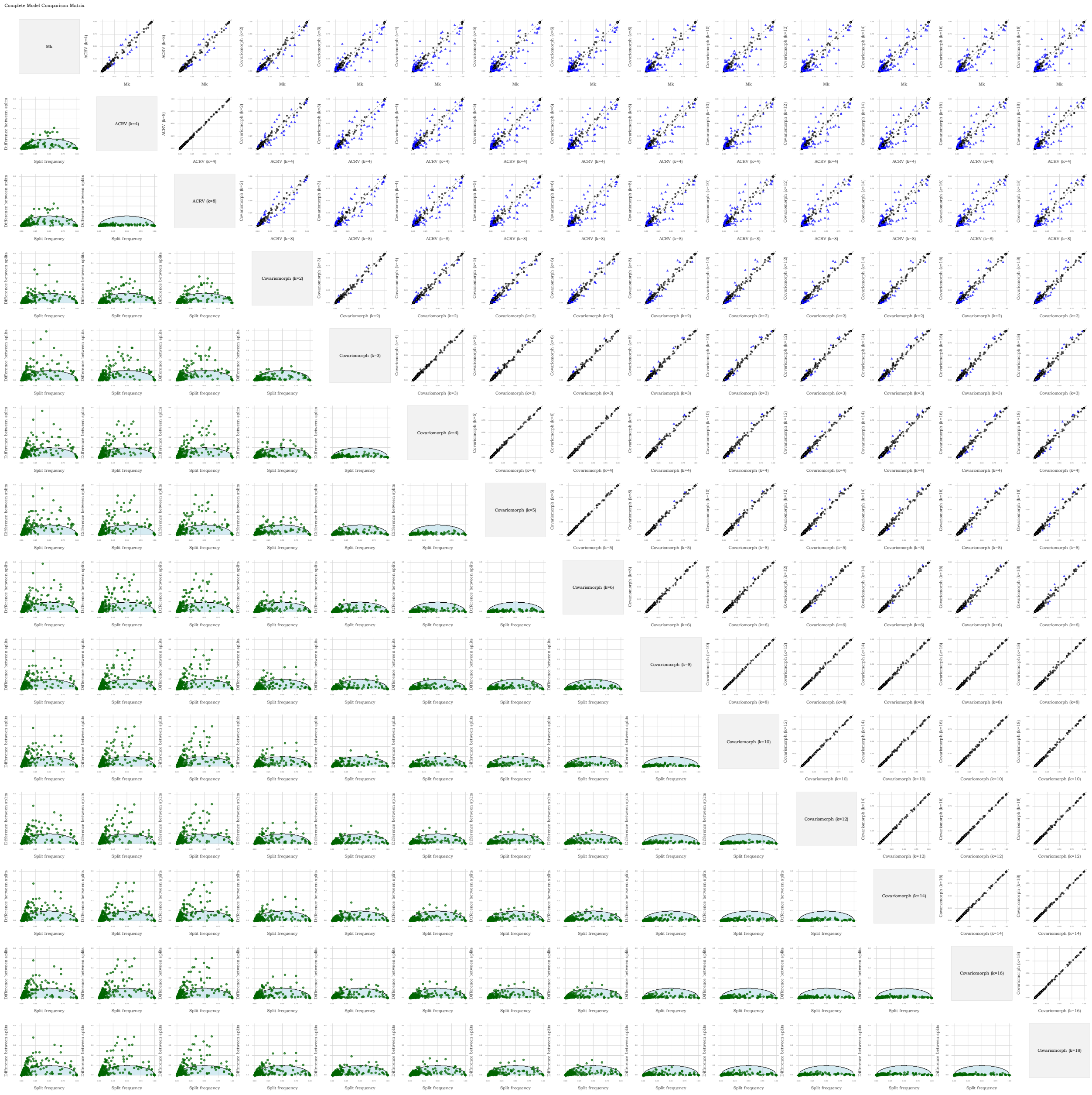

Figure S10: Differences between trees sampled by different models for the Rays dataset. The diagonal gray boxes indicate the models being compared. Each clade is represented by either a circle or a triangle in the plots. **Plots above the diagonal:** These scatter plots show the posterior probabilities of clades, illustrating the level of support each pair of models provides for the same set of potential clades. Circles indicate clades where posterior probabilities do not significantly differ under the expected difference of split frequencies test, whereas triangles represent clades with significantly different posterior probabilities. **Plots below the diagonal** These plots display the differences in split probabilities between model pairs, with the shaded region representing the expected difference of split frequencies for an effective sample size (ESS) of 200. Clades falling within the shaded region pass the expected difference of split frequencies test. If a clade appears above the shaded semicircle, it was sampled with a significantly different probability between the two compared models.

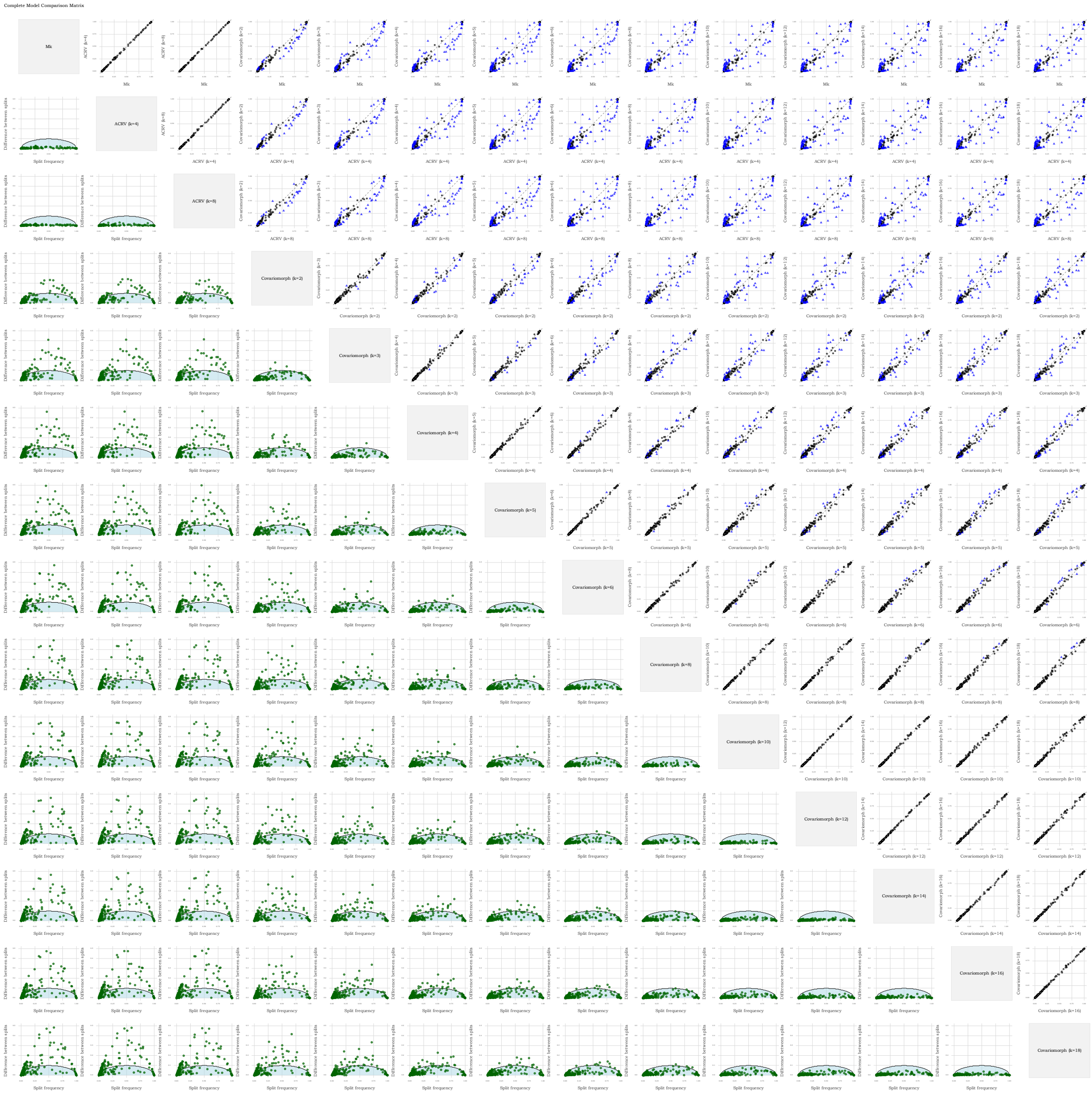

Figure S11: Differences between trees sampled by different models for the Sharks dataset. See Fig. S10 caption for more details on the interpretation.

#### Supplementary References: Data sets used in the main analysis

Abella, J., D. M. Alba, J. M. Robles, A. Valenciano, C. Rotgers, R. Carmona, P. Montoya, and J. Morales. 2012. Kretzoiarctos gen. nov., the oldest member of the giant panda clade. PLoS One 7:e48985.

Abello, M. A. and A. M. Candela. 2020. Paleobiology of argyrolagus (marsupialia, argyrolagidae): an astonishing case of bipedalism among south american mammals. Journal of Mammalian Evolution 27:419–444.

Adrain, J. M. and G. D. Edgecombe. 1997. Silurian encrinurine trilobites from the central Canadian Arctic. Canadian Soc. of Petroleum Geologists.

Ahlberg, P., E. Lukševičs, and E. Mark-Kurik. 2000. A near-tetrapod from the baltic middle devonian. Palaeontology 43:533–548.

Ahlberg, P. E. 1991. A re-examination of sarcopterygian interrelationships, with special reference to the porolepiformes. Zoological Journal of the Linnean Society 103:241–287.

Ahlberg, P. E. and Z. Johanson. 1997. Second tristichopterid (sarcopterygii, osteolepiformes) from the upper devonian of canowindra, new south wales, australia, and phylogeny of the tristichopteridae. Journal of Vertebrate Paleontology 17:653–673.

Allain, R., R. Tykoski, N. Aquesbi, N.-E. Jalil, M. Monbaron, D. Russell, and P. Taquet. 2007. An abelisauroid (dinosauria: Theropoda) from the early jurassic of the high atlas mountains, morocco, and the radiation of ceratosaurs. Journal of Vertebrate Paleontology 27:610–624.

Alvarez, F., R. Jia-Yu, and A. Boucot. 1998. The classification of athyridid brachiopods. Journal of Paleontology 72:827–855.

Amati, L., R. M. Feldmann, and J.-P. Zonneveld. 2004. A new family of triassic lobsters (decapoda: Astacidea) from british columbia and its phylogenetic context. Journal of Paleontology 78:150–168.

Amorim, D. d. S., S. S. Oliveira, D. D. D. do Carmo, and G. C. Ribeiro. 2023. The oldest gondwanan fossil of leiinae (diptera, mycetophilidae): Phylogenetic and evolutionary implications. Cladistics 39:43–57.

Anderson, L. C. and P. D. Roopnarine. 2003. Evolution and phylogenetic relationships of neogene corbulidae (bivalvia; myoidea) of tropical america. Journal of Paleontology 77:1086–1102.

Andruchow-Colombo, A., I. H. Escapa, L. Aagesen, and K. K. Matsunaga. 2023a. In search of lost time: tracing the fossil diversity of podocarpaceae through the ages. Botanical Journal of the Linnean Society 203:315–336.

Andruchow-Colombo, A., G. Rossetto-Harris, T. J. Brodribb, M. A. Gandolfo, and P. Wilf. 2023b. A new fossil acmopyle with accessory transfusion tissue and potential reproductive buds: Direct evidence for ever-wet rainforests in eocene patagonia. American Journal of Botany 110:e16221.

Antoine, P.-O. 2003. Middle miocene elasmotheriine rhinocerotidae from china and mongolia: taxonomic revision and phylogenetic relationships. Zoologica Scripta 32:95–118.

Archibald, J. D., A. O. Averianov, and E. G. Ekdale. 2001. Late cretaceous relatives of rabbits, rodents, and other extant eutherian mammals. Nature 414:62–65.

Argue, D., C. P. Groves, M. S. Lee, and W. L. Jungers. 2017. The affinities of homo floresiensis based on phylogenetic analyses of cranial, dental, and postcranial characters. Journal of Human Evolution 107:107–133.

Argue, D., M. J. Morwood, T. Sutikna, E. W. Saptomo, et al. 2009. Homo floresiensis: a cladistic analysis. Journal of Human Evolution 57:623–639.

Ballen, G. A., J. W. Moreno-Bernal, and C. Jaramillo. 2021. The fossil record of sabre-tooth characins (teleostei: Characiformes: Cynodontinae), their phylogenetic relationships and palaeobiogeographical implications. Journal of Systematic Palaeontology 19:1679–1692.

Baron, M. G. 2019. Pisanosaurus mertii and the triassic ornithischian crisis: could phylogeny offer a solution? Historical Biology 31:967–981.

Barrett, P. Z., S. S. Hopkins, and S. A. Price. 2021. How many sabertooths? reevaluating the number of carnivoran sabretooth lineages with total-evidence bayesian techniques and a novel origin of the miocene nimravidae. Journal of Vertebrate Paleontology 41:e1923523.

Beard, K. C. 1995. The first asian plesiadapoids (mammalia: Primatomorpha). Annals of Carnegie Museum 64:1–33.

Beard, K. C. 2008. The oldest north american primate and mammalian biogeography during the paleocene–eocene thermal maximum. Proceedings of the National Academy of Sciences 105:3815–3818.

Beck, R. M., D. de Vries, M. C. Janiak, I. B. Goodhead, and J. P. Boubli. 2023. Total evidence phylogeny of platyrrhine primates and a comparison of undated and tip-dating approaches. Journal of Human Evolution 174:103293.

Bell, A. and L. M. Chiappe. 2015. Identification of a new hesperornithiform from the cretaceous niobrara chalk and implications for ecologic diversity among early diving birds. PLoS One 10:e0141690.

Benites-Palomino, A., J. Vélez-Juarbe, R. Salas-Gismondi, and M. Urbina. 2019. Scaphokogia totajpe, sp. nov., a new bulky-faced pygmy sperm whale (kogiidae) from the late miocene of peru. Journal of Vertebrate Paleontology 39:e1728538.

Benoit, J., S. Adnet, E. El Mabrouk, H. Khayati, M. Ben Haj Ali, L. Marivaux, G. Merzeraud, S. Merigeaud, M. Vianey-Liaud, and R. Tabuce. 2013. Cranial remain from tunisia provides new clues for the origin and evolution of sirenia (mammalia, afrotheria) in africa. PLoS One 8:e54307.

Benoit, J., S. Adnet, J.-L. Welcomme, and P.-H. Fabre. 2011. New skull of schizodelphis sulcatus gervais, 1861 (mammalia, odontoceti, eurhinodelphinidae) from the lower miocene of pignan (hérault, france) and its implications for systematics of eurhinodelphinidae. Geobios 44:323–334.

Bever, G., T. R. Lyson, D. J. Field, and B.-A. S. Bhullar. 2015. Evolutionary origin of the turtle skull. Nature 525:239–242.

Billet, G. 2010. New observations on the skull of pyrotherium (pyrotheria, mammalia) and new phylogenetic hypotheses on south american ungulates. Journal of Mammalian Evolution 17:21–59.

Billet, G. 2011. Phylogeny of the notoungulata (mammalia) based on cranial and dental characters. Journal of Systematic Palaeontology 9:481–497.

Bloch, J. I., M. T. Silcox, D. M. Boyer, and E. J. Sargis. 2007. New paleocene skeletons and the relationship of plesiadapiforms to crown-clade primates. Proceedings of the National Academy of Sciences 104:1159–1164.

Brownstein, C. 2023. Syngnathoid evolutionary history and the conundrum of fossil misplacement. Integrative Organismal Biology 5:obad011.

Butler, R. J. 2005. The 'fabrosaurid'ornithischian dinosaurs of the upper elliot formation (lower jurassic) of south africa and lesotho. Zoological Journal of the Linnean Society 145:175–218.

Cabreira, S. F., C. L. Schultz, J. S. Bittencourt, M. B. Soares, D. C. Fortier, L. R. Silva, and M. C. Langer. 2011. New stem-sauropodomorph (dinosauria, saurischia) from the triassic of brazil. Naturwissenschaften 98:1035–1040.

Cadena, E. 2015. The first south american sandownid turtle from the lower cretaceous of colombia. PeerJ 3:e1431.

Calamari, Z. T. 2021. Total evidence phylogenetic analysis supports new morphological synapomorphies for bovidae (mammalia, artiodactyla). American Museum Novitates 2021:1–38.

Capobianco, A., S. Zouhri, and M. Friedman. 2025. A long-snouted marine bonytongue (teleostei: Osteoglossidae) from the early eocene of morocco and the phylogenetic affinities of marine osteoglossids. Zoological Journal of the Linnean Society 203:zlae015.

Casali, D. M., A. Boscaini, T. J. Gaudin, and F. A. Perini. 2022. Reassessing the phylogeny and divergence times of sloths (mammalia: Pilosa: Folivora), exploring alternative morphological partitioning and dating models. Zoological Journal of the Linnean Society 196:1505–1551.

Chinnery, B. 2004. Description of prenoceratops pieganensis gen. et sp. nov.(dinosauria: Neoceratopsia) from the two medicine formation of montana. Journal of Vertebrate Paleontology 24:572–590.

Chinnery, B. J. and D. B. Weishampel. 1998. Montanoceratops cerorhynchus (dinosauria: Ceratopsia) and relationships among basal neoceratopsians. Journal of Vertebrate Paleontology 18:569–585.

Clack, J. A., T. J. Challands, T. R. Smithson, and K. Z. Smithson. 2019. Newly recognized famennian lungfishes from east greenland reveal tooth plate diversity and blur the devonian–carboniferous boundary. Papers in Palaeontology 5:261–279.

Claramunt, S. and A. Rinderknecht. 2005. A new fossil furnariid from the pleistocene of uruguay, with remarks on nasal type, cranial kinetics, and relationships of the extinct genus pseudoseisuropsis. The Condor 107:114–127.

Clos, L. M. 1995. A new species of varanus (reptilia: Sauria) from the miocene of kenya. Journal of Vertebrate Paleontology 15:254–267.

Dal Sasso, C., G. Pasini, G. Fleury, and S. Maganuco. 2017. Razanandrongobe sakalavae, a gigantic mesoeucrocodylian from the middle jurassic of madagascar, is the oldest known notosuchian. PeerJ 5:e3481.

Damiani, R. J. and A. M. Yates. 2003. The triassic amphibian thoosuchus yakovlevi and the relationships of the trematosauroidea (temnospondyli: Stereospondyli). Records-Australian Museum 55:331–342.

Darlim, G., M. S. Lee, J. Walter, and M. Rabi. 2022. The impact of molecular data on the phylogenetic position of the putative oldest crown crocodilian and the age of the clade. Biology Letters 18:20210603.

Davis, M. C., N. Shubin, and E. B. Daeschler. 2004. A new specimen of sauripterus taylori (sarcopterygii, osteichthyes) from the famennian catskill formation of north america. Journal of Vertebrate Paleontology 24:26–40.

Davis, M. P., G. Arratia, and T. M. Kaiser. 2013. The first fossil shellear and its implications for the evolution and divergence of the kneriidae (teleostei: Gonorynchiformes). Mesozoic Fishes 5:325–362.

Delcourt, R. and O. N. Grillo. 2018. Tyrannosauroids from the southern hemisphere: implications for biogeography, evolution, and taxonomy. Palaeogeography, Palaeoclimatology, Palaeoecology 511:379–387.

Dembo, M., D. Radovčić, H. M. Garvin, M. F. Laird, L. Schroeder, J. E. Scott, J. Brophy, R. R. Ackermann, C. M. Musiba, D. J. de Ruiter, et al. 2016. The evolutionary relationships and age of homo naledi: An assessment using dated bayesian phylogenetic methods. Journal of Human Evolution 97:17–26.

Doran Brownstein, C., L. Yang, M. Friedman, and T. J. Near. 2023. Phylogenomics of the ancient and species-depauperate gars tracks 150 million years of continental fragmentation in the northern hemisphere. Systematic Biology 72:213–227.

D'Emic, M. D. 2013. Revision of the sauropod dinosaurs of the lower cretaceous trinity group, southern usa, with the description of a new genus. Journal of Systematic Palaeontology 11:707–726.

Eddy, D. R. and J. A. Clarke. 2011. New information on the cranial anatomy of acrocanthosaurus atokensis and its implications for the phylogeny of allosauroidea (dinosauria: Theropoda). PloS One 6:e17932.

Everett, C. J., T. A. Deméré, and A. R. Wyss. 2023. A new species of pinnarctidion from the pysht formation of washington state (usa) and a phylogenetic analysis of basal pan-pinnipeds (eutheria, carnivora). Journal of Vertebrate Paleontology 42.

Evers, S. W., K. E. Chapelle, and W. G. Joyce. 2023. Cranial and mandibular anatomy of plastomenus thomasii and a new time-tree of trionychid evolution. Swiss Journal of Palaeontology 142:1.

Ezcurra, M. D. and G. Cuny. 2007. The coelophysoid lophostropheus airelensis, gen. nov.: a review of the systematics of "liliensternus" airelensis from the triassic–jurassic outcrops of normandy (france). Journal of Vertebrate Paleontology 27:73–86.

Fanti, F., G. L. Conte, L. Angelicola, and A. Cau. 2016. Why so many dipnoans? a multidisciplinary approach on the lower cretaceous lungfish record from tunisia. Palaeogeography, Palaeoclimatology, Palaeoecology 449:255–265.

Ferratges, F. A., J. Luque, J. L. Domínguez, À. Ossó, M. Aurell, and S. Zamora. 2023. The oldest dairoidid crab (decapoda, brachyura, parthenopoidea) from the eocene of spain. Papers in Palaeontology 9:e1494.

Field, D. J., J. Benito, A. Chen, J. W. Jagt, and D. T. Ksepka. 2020. Late cretaceous neornithine from europe illuminates the origins of crown birds. Nature 579:397–401.

Ford, S. M. 1994. Primitive platyrrhines? perspectives on anthropoid origins from platyrrhine, parapithecid, and preanthropoid postcrania. Anthropoid Origins Pages 595–673.

Friis, E. M., K. R. Pedersen, M. von Balthazar, G. W. Grimm, and P. R. Crane. 2009. Monetianthus mirus gen. et sp. nov., a nymphaealean flower from the early cretaceous of portugal. International Journal of Plant Sciences 170:1086–1101.

Gasulla, J. M., F. Escaso, I. Narváez, F. Ortega, and J. L. Sanz. 2015. A new sail-backed styracosternan (dinosauria: Ornithopoda) from the early cretaceous of morella, spain. PLoS One 10:e0144167.

Gebo, D. L., M. Dagosto, K. C. Beard, and T. Qi. 2001. Middle eocene primate tarsals from china: implications for haplorhine evolution. American Journal of Physical Anthropology: The Official Publication of the American Association of Physical Anthropologists 116:83–107.

Geroto, C. F. C. and R. J. Bertini. 2019. New material of pepesuchus (crocodyliformes; mesoeucrocodylia) from the bauru group: implications about its phylogeny and the age of the adamantina formation. Zoological Journal of the Linnean Society 185:312–334.

Gonçalves, R. B., O. M. De Meira, and B. B. Rosa. 2022. Total-evidence dating and morphological partitioning: a novel approach to understand the phylogeny and biogeography of augochlorine bees (hymenoptera: Apoidea). Zoological Journal of the Linnean Society 195:1390–1406.

Groh, S. S., P. Upchurch, P. M. Barrett, and J. J. Day. 2020. The phylogenetic relationships of neosuchian crocodiles and their implications for the convergent evolution of the longirostrine condition. Zoological Journal of the Linnean Society 188:473–506.

Harper, E. M., E. A. Hide, and B. Morton. 2000. Relationships between the extant anomalodesmata: a cladistic test. *in* The Evolutionary Biology of the Bivalvia. Geological Society of London.

Heckeberg, N. S. 2020. The systematics of the cervidae: a total evidence approach. PeerJ 8:e8114.

Hendrickx, C., L. C. Gaetano, J. N. Choiniere, H. Mocke, and F. Abdala. 2020. A new traversodontid cynodont with a peculiar postcanine dentition from the middle/late triassic of namibia and dental evolution in basal gomphodonts. Journal of Systematic Palaeontology 18:1669–1706.

Holbrook, L. T. 1999. The phylogeny and classification of tapiromorph perissodactyls (mammalia). Cladistics 15:331–350.

Holley, J. A., J. Sterli, and N. G. Basso. 2020. Dating the origin and diversification of pan-chelidae (testudines, pleurodira) under multiple molecular clock approaches. Contributions to Zoology 89:146–174.

Jin, P., M. Zhang, B. Du, A. Li, and B. Sun. 2023. A new species of pararaucaria from the lower cretaceous of shandong province (eastern china): Insights into the evolution of the cheirolepidiaceae cone. Cretaceous Research 146:105475.

Johanson, Z. 2004. Late devonian sarcopterygian fishes from eastern gondwana (australia and antarctica) and their importance in phylogeny and biogeography. Recent Advances in the Origin and Early Radiation of Vertebrates Pages 287–308.

Jouault, C., A. Maréchal, F. L. Condamine, B. Wang, A. Nel, F. Legendre, and V. Perrichot. 2022. Including fossils in phylogeny: a glimpse into the evolution of the superfamily evanioidea (hymenoptera: Apocrita) under tip-dating and the fossilized birth–death process. Zoological Journal of the Linnean Society 194:1396–1423.

Karasawa, H. and H. Kato. 2003. The family goneplacidae macleay, 1838 (crustacea: Decapoda: Brachyura): systematics, phylogeny, and fossil records. Paleontological Research 7:129–151.

Kay, R. F., J. Thewissen, and A. D. Yoder. 1992. Cranial anatomy of ignacius graybullianus and the affinities of the plesiadapiformes. American Journal of Physical Anthropology 89:477–498.

Kay, R. F. and B. A. Williams. 1994. Dental evidence for anthropoid origins. Anthropoid Origins Pages 361–445.

Kim, J., G. Cassis, and S. Jung. 2023. Phylogenetic analysis of the predatory plant bug subfamily deraeocorinae (hemiptera: Heteroptera: Miridae) based on molecular and morphological data. Zoological Journal of the Linnean Society 197:246–266.

Laloy, F., J.-C. Rage, S. E. Evans, R. Boistel, N. Lenoir, and M. Laurin. 2013. A re-interpretation of the eocene anuran thaumastosaurus based on microct examination of a 'mummified' specimen. PLoS One 8:e74874.

Langer, M. C., A. D. Rincón, J. Ramezani, A. Solórzano, and O. W. Rauhut. 2014. New dinosaur (theropoda, stem-averostra) from the earliest jurassic of the la quinta formation, venezuelan andes. Royal Society Open Science 1:140184.

Larsson, H. C. and H.-D. Sues. 2007. Cranial osteology and phylogenetic relationships of hamadasuchus rebouli (crocodyliformes: Mesoeucrocodylia) from the cretaceous of morocco. Zoological Journal of the Linnean Society 149:533–567.

Lee, K., J. Feinstein, J. Cracraft, and D. Mindell. 1997. The phylogeny of ratite birds: resolving conflicts between molecular and morphological data sets. Avian Molecular Evolution and Systematics Pages 173–211.

Lee, M. S. 1995. Historical burden in systematics and the interrelationships of 'parareptiles'. Biological Reviews 70:459–547.

Lee, M. S. and J. D. Scanlon. 2002. The cretaceous marine squamate mesoleptos and the origin of snakes. Bulletin of the Natural History Museum: Zoology Series 68:131–142.

Li, Y.-D., S. Yamamoto, A. F. Newton, and C.-Y. Cai. 2023. Kekveus brevisulcatus sp. nov., a new featherwing beetle from mid-cretaceous amber of northern myanmar (coleoptera: Ptiliidae). PeerJ 11:e15306.

Liu, J., L.-J. Zhao, C. Li, and T. He. 2013. Osteology of concavispina biseridens (reptilia, thalattosauria) from the xiaowa formation (carnian), guanling, guizhou, china. Journal of Paleontology 87:341–350.

Lloyd, G. T., S. C. Wang, and S. L. Brusatte. 2012. Identifying heterogeneity in rates of morphological evolution: discrete character change in the evolution of lungfish (sarcopterygii; dipnoi). Evolution 66:330–348.

Long-Fox, B. L. 2022. Phylogenetic controls on shell morphology in the chemosymbiotic Lucinidae (Bivalvia). South Dakota School of Mines and Technology.

Longrich, N. 2006. An ornithurine bird from the late cretaceous of alberta, canada. Canadian Journal of Earth Sciences 43:1–7.

Lu, J., S. Giles, M. Friedman, and M. Zhu. 2017. A new stem sarcopterygian illuminates patterns of character evolution in early bony fishes. Nature Communications 8:1932.

Lü, J., G. Li, M. Kundrát, Y.-N. Lee, Z. Sun, Y. Kobayashi, C. Shen, F. Teng, and H. Liu. 2017. High diversity of the ganzhou oviraptorid fauna increased by a new "cassowary-like" crested species. Scientific Reports 7:6393.

Luckett, W. P. 1993. Developmental evidence from the fetal membranes for assessing archontan relationships. Pages 149–186 *in* Primates and their relatives in phylogenetic perspective. Springer.

Lundberg, J. G. 1992. The phylogeny of ictalurid catfishes: a synthesis of recent work. Systematics, Historical Ecology, and North American Freshwater Fishes 392:420.

Magalhaes, I. L. and M. J. Ramírez. 2022. Phylogeny and biogeography of the ancient spider family filistatidae (araneae) is consistent both with long-distance dispersal and vicariance following continental drift. Cladistics 38:538–562.

Mallon, J. C., C. J. Ott, P. L. Larson, E. M. Iuliano, and D. C. Evans. 2016. Spiclypeus shipporum gen. et sp. nov., a boldly audacious new chasmosaurine ceratopsid (dinosauria: Ornithischia) from the judith river formation (upper cretaceous: Campanian) of montana, usa. PloS One 11:e0154218.

Mao, F., C. Zhang, C. Liu, and J. Meng. 2021. Fossoriality and evolutionary development in two cretaceous mammaliamorphs. Nature 592:577–582.

Marivaux, L. 2006. The eosimiid and amphipithecid primates (anthropoidea) from the oligocene of the bugti hills (balochistan, pakistan): new insight into early higher primate evolution in south asia. Palaeovertebrata .

Marivaux, L., M. Adaci, M. Bensalah, H. G. Rodrigues, L. Hautier, M. Mahboubi, F. Mebrouk, R. Tabuce, and M. Vianey-Liaud. 2011. Zegdoumyidae (rodentia, mammalia), stem anomaluroid rodents from the early to middle eocene of algeria (gour lazib, western sahara): new dental evidence. Journal of Systematic Palaeontology 9:563–588.

Marivaux, L., A. Ramdarshan, E. M. Essid, W. Marzougui, H. K. Ammar, R. Lebrun, B. Marandat, G. Merzeraud, R. Tabuce, and M. Vianey-Liaud. 2013. Djebelemur, a tiny pre-tooth-combed primate from the eocene of tunisia: a glimpse into the origin of crown strepsirhines. PLoS One 8:e80778.

Marramà, G., E. Villalobos-Segura, R. Zorzin, J. Kriwet, and G. Carnevale. 2023. The evolutionary origin of the durophagous pelagic stingray ecomorph. Palaeontology 66:e12669.

Martínez, R. N. and C. Apaldetti. 2017. A late norian—rhaetian coelophysid neotheropod (dinosauria, saurischia) from the quebrada del barro formation, northwestern argentina. Ameghiniana 54:488–505.

Masters, J. C. and D. J. Brothers. 2002. Lack of congruence between morphological and molecular data in reconstructing the phylogeny of the galagonidae. American Journal of Physical Anthropology: The Official Publication of the American Association of Physical Anthropologists 117:79–93.

Mateus, O., E. Puértolas-Pascual, and P. M. Callapez. 2019. A new eusuchian crocodylomorph from the cenomanian (late cretaceous) of portugal reveals novel implications on the origin of crocodylia. Zoological Journal of the Linnean Society 186:501–528.

Mauricio, G. N., J. I. Areta, M. R. Bornschein, and R. E. Reis. 2012. Morphology-based phylogenetic analysis and classification of the family rhinocryptidae (aves: Passeriformes). Zoological Journal of the Linnean Society 166:377–432.

May, M. R., D. L. Contreras, M. A. Sundue, N. S. Nagalingum, C. V. Looy, and C. J. Rothfels. 2021. Inferring the total-evidence timescale of marattialean fern evolution in the face of model sensitivity. Systematic Biology 70:1232–1255.

Mayr, G. 2016. Osteology and phylogenetic affinities of the middle eocene north american bathornis grallator—one of the best represented, albeit least known paleogene cariamiform birds (seriemas and allies). Journal of Paleontology 90:357–374.

Mayr, G., V. L. De Pietri, R. P. Scofield, and T. Smith. 2019. A fossil heron from the early oligocene of belgium: the earliest temporally well-constrained record of the ardeidae. Ibis 161:79–90.

Mayr, G. and A. C. Kitchener. 2022. Oldest fossil loon documents a pronounced ecomorphological shift in the evolution of gaviiform birds. Zoological Journal of the Linnean Society 196:1431–1450.

Mayr, G. and A. C. Kitchener. 2023. A new fossil from the london clay documents the convergent origin of a "mousebird-like" tarsometatarsus in an early eocene near-passerine bird. Acta Palaeontologica Polonica 68:1–11.

Mihlbachler, M. C., S. G. Lucas, R. J. Emry, and B. Bayshashov. 2004. A new brontothere (brontotheriidae, perissodactyla, mammalia) from the eocene of the ily basin of kazakstan and a phylogeny of asian "horned" brontotheres. American Museum Novitates 2004:1–43.

Mongle, C. S., D. S. Strait, and F. E. Grine. 2023. An updated analysis of hominin phylogeny with an emphasis on re-evaluating the phylogenetic relationships of australopithecus sediba. Journal of Human Evolution 175:103311.

Moon, J., J.-B. Caron, and J. Moysiuk. 2023. A macroscopic free-swimming medusa from the middle cambrian burgess shale. Proceedings of the Royal Society B 290:20222490.

Morales, M. and M. A. Shishkin. 2002. A re-assessment of parotosuchus africanus (broom), a capitosauroid temnospondyl amphibian from the triassic of south africa. Journal of Vertebrate Paleontology 22:1–11.

Murray, A. M., L. E. Nelson, and D. B. Brinkman. 2023. A new sturgeon from the upper cretaceous horseshoe canyon formation in central alberta, canada. Journal of Vertebrate Paleontology 43:e2232846.

Norell, M. A. 1989. The higher level relationships of the extant crocodylia. Journal of Herpetology Pages 325–335.

Otero, R. A., J. P. O'Gorman, N. Hiller, F. R. O'Keefe, and R. E. Fordyce. 2016. Alexandronectes zealandiensis gen. et sp. nov., a new aristonectine plesiosaur from the lower maastrichtian of new zealand. Journal of Vertebrate Paleontology 36:e1054494.

O'Meara, R. N. and R. S. Thompson. 2014. Were there miocene meridiolestidans? assessing the phylogenetic placement of necrolestes patagonensis and the presence of a 40 million year meridiolestidan ghost lineage. Journal of Mammalian Evolution 21:271–284.

Parker, W. G. 2018. Anatomical notes and discussion of the first described aetosaur stagonolepis robertsoni (archosauria: Suchia) from the upper triassic of europe, and the use of plesiomorphies in aetosaur biochronology. PeerJ 6:e5455.

Peecook, B. R., C. A. Sidor, S. J. Nesbitt, R. M. Smith, J. S. Steyer, and K. D. Angielczyk. 2013. A new silesaurid from the upper ntawere formation of zambia (middle triassic) demonstrates the rapid diversification of silesauridae (avemetatarsalia, dinosauriformes). Journal of Vertebrate Paleontology 33:1127–1137.

Penkalski, P. 2018. Revised systematics of the armoured dinosaur euoplocephalus and its allies. Neues Jahrbuch für Geologie und Paläontologie-Abhandlungen 287:261–306.

Ramsköld, L. and L. Werdelin. 1991. The phylogeny and evolution of some phacopid trilobites. Cladistics 7:29–74.

Randle, E. and R. S. Sansom. 2017. Phylogenetic relationships of the 'higher heterostracans'(heterostraci: Pteraspidiformes and cyathaspididae), extinct jawless vertebrates. Zoological Journal of the Linnean Society 181:910–926.

Rangel, C. C., L. M. Carneiro, M. F. Tejedor, L. P. Bergqvist, and É. V. Oliveira. 2023. A reassessment of nemolestes (mammalia, metatheria): Systematics and evolutionary implications for sparassodonta. Journal of Mammalian Evolution 30:535–559.

Rauhut, O. W., A. M. Heyng, A. López-Arbarello, and A. Hecker. 2012. A new rhynchocephalian from the late jurassic of germany with a dentition that is unique amongst tetrapods. PLoS One 7:e46839.

Ristevski, J., M. T. Young, M. B. De Andrade, and A. K. Hastings. 2018. A new species of anteophthalmosuchus (crocodylomorpha, goniopholididae) from the lower cretaceous of the isle of wight, united kingdom, and a review of the genus. Cretaceous Research 84:340–383.

Rode, A. L. and L. E. Babcock. 2003. Phylogeny of fossil and extant freshwater crayfish and some closely related nephropid lobsters. Journal of Crustacean Biology 23:418–435.

Rothwell, G. W. and K. C. Nixon. 2006. How does the inclusion of fossil data change our conclusions about the phylogenetic history of euphyllophytes? International Journal of Plant Sciences 167:737–749.

Royo-Torres, R., C. Fuentes, M. Meijide, F. Meijide-Fuentes, and M. Meijide-Fuentes. 2017. A new brachiosauridae sauropod dinosaur from the lower cretaceous of europe (soria province, spain). Cretaceous Research 80:38–55.

Rule, J. P., J. W. Adams, and E. M. Fitzgerald. 2021. Early monk seals (monachinae: Monachini) from the late miocene–early pliocene of australia. Journal of Systematic Palaeontology 19:441–459.

Sarigul, V., F. Agnolin, and S. Chatterjee. 2018. Description of a multitaxic bone assemblage from the upper triassic post quarry of texas (dockum group), including a new small basal dinosauriform taxon. Historia Natural 8:5–24.

Scanlon, J. D. and M. S. Lee. 2000. The pleistocene serpent wonambi and the early evolution of snakes. Nature 403:416–420.

Scarpetta, S. G. 2024. A palaeogene stem crotaphytid (aciprion formosum) and the phylogenetic affinities of early fossil pleurodontan iguanians. Royal Society Open Science 11:221139.

Scarpetta, S. G., D. T. Ledesma, and C. J. Bell. 2021. A new extinct species of alligator lizard (squamata: Elgaria) and an expanded perspective on the osteology and phylogeny of gerrhonotinae. BMC Ecology and Evolution 21:1–58.

Schaefer, S. A. 1987. Osteology of Hypostomus Plecostomus (linnaeus): With a Phylogenetic Analysis of the Loricariid Subfamilies (Pisces, Siluroidei). Natural History Museum of Los Angeles County.

Schoch, R. R. and A. R. Milner. 2008. The intrarelationships and evolutionary history of the temnospondyl family branchiosauridae. Journal of Systematic Palaeontology 6:409–431.

Shirai, S. 1996. Phylogenetic interrelationships of neoselachians (chondrichthyes: Euselachii). Interrelationships of Fishes Pages 9–34.

Silcox, M. T., D. W. Krause, M. C. Maas, and R. C. Fox. 2001. New specimens of elphidotarsius russelli (mammalia,? primates, carpolestidae) and a revision of plesiadapoid relationships. Journal of Vertebrate Paleontology 21:132–152.

Simmons, N. B. and T. M. Conway. 2001. Phylogenetic relationships of mormoopid bats (chiroptera: Mormoopidae) based on morphological data. Bulletin of the American Museum of Natural History 2001:1–100.

Simões, T. R., O. Vernygora, M. W. Caldwell, and S. E. Pierce. 2020. Megaevolutionary dynamics and the timing of evolutionary innovation in reptiles. Nature Communications 11:3322.

Smith, K. T. and A. Scanferla. 2021. A nearly complete skeleton of the oldest definitive erycine boid (messel, germany). Geodiversitas 43:1–24.

Tanaka, Y. and M. Watanabe. 2019. An early and new member of balaenopteridae from the upper miocene of hokkaido, japan. Journal of Systematic Palaeontology 17:1417–1431.

Tattersall, I. 1993. Speciation and morphological differentiation in the genus lemur. Pages 163–176 *in* Species, Species Concepts and Primate Evolution. Springer.

Thomson, S. and A. Georges. 2009. Myuchelys gen. nov.—a new genus for elseya latisternum and related forms of australian freshwater turtle (testudines: Pleurodira: Chelidae). Zootaxa 2053:32–42.

Travouillon, K., S. Hand, M. Archer, and K. Black. 2014. Earliest modern bandicoot and bilby (marsupialia, peramelidae and thylacomyidae) from the miocene of the riversleigh world heritage area, northwestern queensland, australia. Journal of Vertebrate Paleontology 34:375–382.

Valenti, P., E. Vlachos, C. Kehlmaier, U. Fritz, G. L. Georgalis, À. H. Luján, R. Miccichè, L. Sineo, and M. Delfino. 2022. The last of the large-sized tortoises of the mediterranean islands. Zoological Journal of the Linnean Society 196:1704–1717.

Vigliotta, T. R. 2008. A phylogenetic study of the african catfish family mochokidae (osteichthyes, ostariophysi, siluriformes), with a key to genera. Proceedings of the Academy of Natural Sciences of Philadelphia 157:73–136.

Vlachos, E. and M. Rabi. 2018. Total evidence analysis and body size evolution of extant and extinct tortoises (testudines: Cryptodira: Pan-testudinidae). Cladistics 34:652–683.

Walker, Z., G. W. Rothwell, and R. A. Stockey. 2023. Fossil evidence for sporeling development of a mesozoic osmundaceous fern. American Journal of Botany 110:e16210.

Wang, Y., W. A. Clemens, Y. Hu, and C. Li. 1998. A probable pseudo-tribosphenic upper molar from the late jurassic of china and the early radiation of the holotheria. Journal of Vertebrate Paleontology 18:777–787.

Westrop, S. R., R. Ludvigsen, and C. H. Kindle. 1996. Marjuman (cambrian) agnostoid trilobites of the cow head group, western newfoundland. Journal of Paleontology 70:804–829.

Wiersma, J. P. and R. B. Irmis. 2018. A new southern laramidian ankylosaurid, akainacephalus johnsoni gen. et sp. nov., from the upper campanian kaiparowits formation of southern utah, usa. PeerJ 6:e5016.

Williams, B. A. and H. H. Covert. 1994. New early eocene anaptomorphine primate (omomyidae) from the washakie basin, wyoming, with comments on the phylogeny and paleobiology of anaptomorphines. American Journal of Physical Anthropology 93:323–340.

Williamson, T. E., S. L. Brusatte, R. Secord, and S. Shelley. 2016. A new taeniolabidoid multituberculate (mammalia) from the middle puercan of the nacimiento formation, new mexico, and a revision of taeniolabidoid systematics and phylogeny. Zoological Journal of the Linnean Society 177:183–208.

Xiang, X., K. Xiang, R. D. C. Ortiz, F. Jabbour, and W. Wang. 2019. Integrating palaeontological and molecular data uncovers multiple ancient and recent dispersals in the pantropical hamamelidaceae. Journal of Biogeography 46:2622–2631.

Yates, A. M., J. Ristevski, and S. W. Salisbury. 2023. The last baru (crocodylia, mekosuchinae): a new species of 'cleaver-headed crocodile' from central australia and the turnover of crocodylians during the late miocene in australia. Papers in Palaeontology 9:e1523.

Zaher, H., D. M. Mohabey, F. G. Grazziotin, and J. A. Wilson Mantilla. 2023. The skull of sanajeh indicus, a cretaceous snake with an upper temporal bar, and the origin of ophidian wide-gaped feeding. Zoological Journal of the Linnean Society 197:656–697.

Zanata, A. M. and R. P. Vari. 2005. The family alestidae (ostariophysi, characiformes): a phylogenetic analysis of a trans-atlantic clade. Zoological Journal of the Linnean Society 145:1–144.
